# Dietary Pyruvate Targets Cytosolic Phospholipase A2 to Mitigate Inflammation and Obesity in Mice

**DOI:** 10.1101/2023.09.07.556702

**Authors:** Sadaf Hasan, Nabil Ghani, Xiangli Zhao, Julia Good, Amanda Huang, Hailey Lynn Wrona, Jody Liu, Chuan-ju Liu

## Abstract

Obesity has a multifactorial etiology and is known to be a state of chronic low-grade inflammation, known as meta-inflammation. This state is associated with the development of metabolic disorders such as glucose intolerance and nonalcoholic fatty liver disease. Pyruvate is a glycolytic metabolite and a crucial node in various metabolic pathways. However, its role and molecular mechanism in obesity and associated complications are obscure. In this study, we reported that pyruvate substantially inhibited adipogenic differentiation *in vitro* and its administration significantly prevented HFD-induced weight gain, white adipose tissue inflammation, and metabolic dysregulation. To identify the target proteins of pyruvate, drug affinity responsive target stability was employed with proteomics, cellular thermal shift assay, and isothermal drug response to detect the interactions between pyruvate and its molecular targets. Consequently, we identified cytosolic phospholipase A2 (cPLA2) as a novel molecular target of pyruvate and demonstrated that pyruvate restrained diet-induced obesity, white adipose tissue inflammation, and hepatic steatosis in a cPLA2-dependent manner. Studies with global ablation of cPLA2 in mice showed that the protective effects of pyruvate were largely abrogated, confirming the importance of pyruvate/cPLA2 interaction in pyruvate attenuation of inflammation and obesity. Overall, our study not only establishes pyruvate as an antagonist of cPLA2 signaling and a potential therapeutic option for obesity, but it also sheds light on the mechanism of its action. Pyruvate’s prior clinical use indicates that it can be considered a safe and viable alternative for obesity, whether consumed as a dietary supplement or as part of a regular diet.

## Introduction

Obesity is a growing epidemic that originates from excessive caloric intake and a positive energy balance [1]. As a complex condition, its pathophysiology cannot be attributed only to energy imbalances between intake and expenditure [2]. Nevertheless, it is well-established that obesity is associated with an increase in inflammatory markers, leading to persistent chronic low-grade inflammation, which results in altered metabolism, termed meta-inflammation [3]. Thus, recently its etiology has been reconsidered as inflammatory diseases driven by metabolic dysregulation [4]. This meta-inflammation is believed to be a crucial factor in the manifestation of obesity-related comorbidities such as nonalcoholic fatty liver disease (NAFLD), cardiovascular diseases, type 2 diabetes mellitus, and neurodegenerative diseases [5].

At the cellular level, obesity is characterized by an increase in adipogenesis [6]. Adipogenesis refers to the process by which preadipocytes differentiate into adipocytes capable of storing excess energy in the form of fat [7], making adipose tissue a vital metabolic organ and a critical player in the regulation of whole-body energy homeostasis [8]. Obesity is intimately correlated with adipocyte hypertrophy (cell size increase) and/or hyperplasia (cell number increase), which promote lipid accumulation and enhance immune cell activity in adipose tissue, triggering repercussions in tissue physiology.

By convention, adipose tissues (AT) are classified into two categories, brown adipose tissue (BAT) and white adipose tissue. Inherently, white and brown adipocytes differ in their morphology, resulting in their divergent functions [9]. BAT modulates body temperature primarily through non-shivering thermogenesis, the dissipation of chemical energy to generate heat [10]. However, WAT plays a protagonist in the pathogenesis of metabolic disorders associated with obesity.

WAT is the primary location of inflammatory response in obesity [11]. In response to weight gain and adiposity, it undergoes a phenotypic switch that is characterized by the presence of inflamed, dysfunctional adipocytes in association with immune cell infiltration, mainly macrophages. The adipocyte-macrophage crosstalk perpetuates a vicious cycle of macrophage recruitment and NF-κB-dependent expression of proinflammatory cytokines such as interleukin– 1β (IL−1β), interleukin-6 (IL-6), and tumor necrosis factor-α (TNFα). This axis in turn disrupts the optimal function of the AT itself and of other remote organs [12].

In the adipose tissue microenvironment, macrophages are remarkably heterogeneous and highly adaptable. There are two main subpopulations involved in obesity-related inflammation viz., M1 or classically activated (proinflammatory) and M2 or alternatively activated (anti-inflammatory) macrophages. In lean adipose tissue, M2 macrophages are predominant and participate actively in suppressing the inflammatory response preserving adipose tissue homeostasis [13]. However, in an obese state, AT is infiltrated with proinflammatory M1 macrophages, exacerbating both local and systemic inflammation. Hence, impeding macrophage infiltration and skewing the phenotypic profile of M1/M2 macrophages in WAT will be an effective treatment strategy for obesity-related inflammation.

Pyruvate is a product of cytosolic glycolysis and a crucial node for multiple metabolic pathways. Besides serving as an energy-bearing metabolite, it acts as an endogenous antioxidant and anti-inflammatory molecule [14]. On the other hand, exogenous pyruvate which is incorporated into cells via monocarboxylate transporters (MCTs), has been reported to ameliorate retinopathy, nephropathy in streptozotocin-induced diabetic mice, and hyperglycemia [15]. Moreover, it is described to improve cardiac function after coronary ischemia and reperfusion as well as in critical medical conditions, such as severe sepsis [16].

Interestingly, there is limited clinical evidence, dating back more than two decades, that the administration of pyruvate decreases body weight in obese individuals [17]. However, the target and the underlying mechanism of pyruvate remains unknown. This led us to explore the molecular target and mechanism of action of pyruvate in high fat diet (HFD) induced obesity. To the best of our knowledge, this is the first study to demonstrate that pyruvate inhibits adipogenesis and protects against HFD-induced weight gain by targeting cytosolic phospholipase A2 (cPLA2), which is known to play a critical role in adipocytic differentiation [18] and in chronic inflammation [19].

## Results

### Pyruvate inhibits adipogenic differentiation

RNA-seq analysis revealed that pyruvate administration downregulated a plethora of genes in a systemic model of inflammation including genes playing a role in inflammation and/or obesity (**Fig. 1A**). The volcano plot also confirmed significant downregulation of various genes in inflammation and obesity including IL-6, IL-1Β, CCL2, PPARg, and Fabp4 (**Fig. 1B**). We incubated primary murine preadipocytes with various concentrations of pyruvate during 8-day-long differentiation. The pyruvate concentration ranged from 0.02 to 10 mM, where 1 mM is the standard concentration that is used in cell culture media and 0.02 mM corresponds to physiological plasma levels in humans. qPCR analyses indicated that the expression of the adipogenic transcriptional markers (PPARγ and CEBPα) as well as of an adipocyte-related gene (Fabp4) was suppressed by pyruvate in a V-shape trend indicating a concentration-dependent inhibition between 2 mM and 4 mM. Pyruvate at 8 mM also showed significant suppression of the expression levels of the markers but 4 mM exhibited the most significant effect. **(Fig 1C-E**). Further assessments of the time-induced effects from day 0 – day 8 were performed using 4 mM pyruvate. This qPCR analysis showed that mRNA levels of PPARγ, CEBPα, and Fabp4 were significantly suppressed during adipogenic induction by pyruvate **(Fig. 1F-H**). As a crucial parameter of toxicity assays, we also evaluated pyruvate’s effect on cell viability. An MTT assay, using a range of pyruvate concentrations, for seven days over three different time points, demonstrated that pyruvate had an indistinguishable effect on cell viability (**Fig. S1A**).

**Figure 1:**
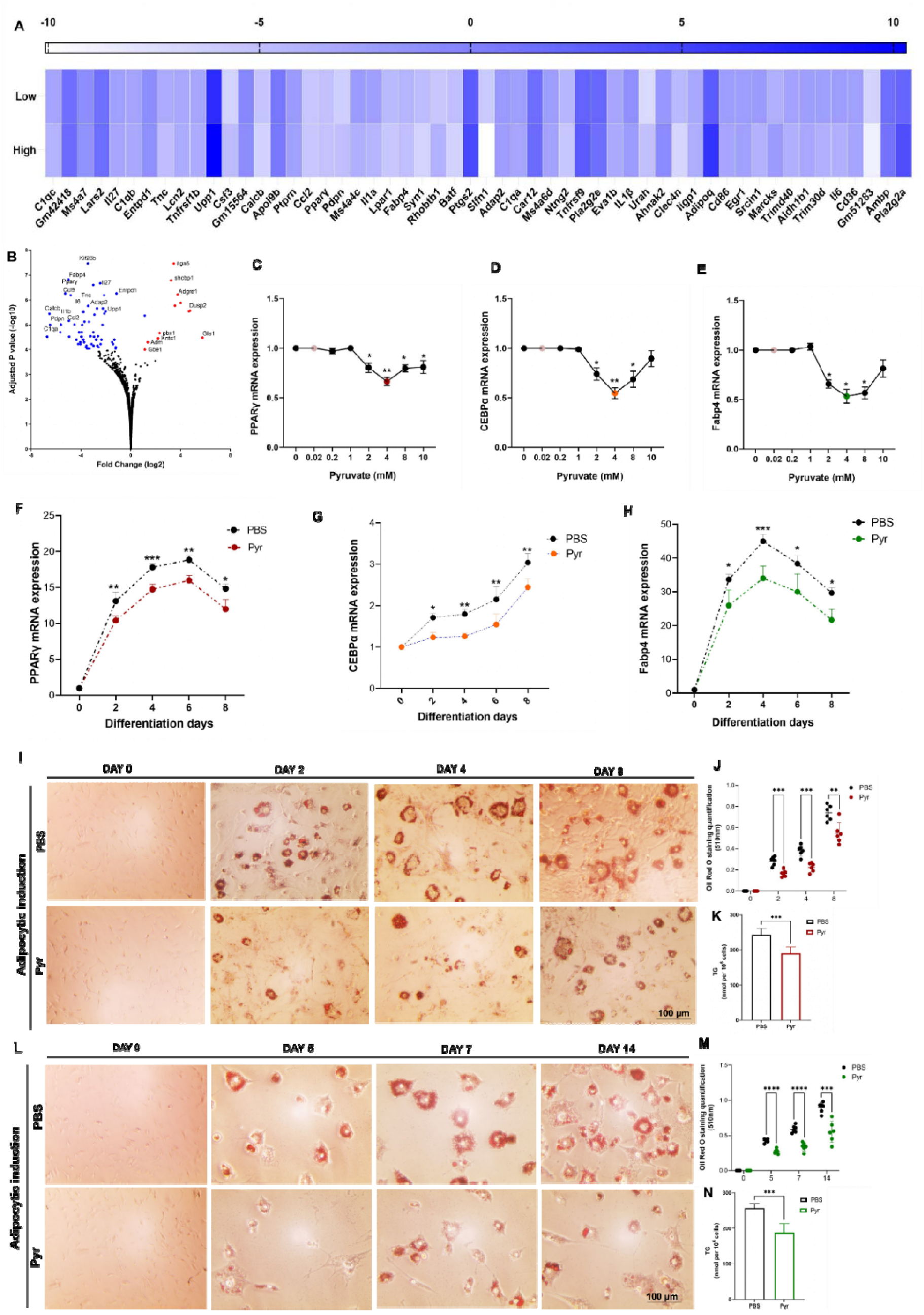
Effect of pyruvate on adipogenic differentiation in primary cells. **A**. Heatmap of gene expression levels from RNA seq analysis of bmMSCs isolated from C57BL/6 mice treated with or without low (2 mM) or high (4 mM) concentrations of pyruvate, in the presence of TNFα for 24 hours. **B**. Volcano plot showing statistical significance (P-value) vs fold change for differentially expressed genes (DEGs) of bmMSCs treated with pyruvate in the presence (4 mM) in the presence of TNFα for 24 hours. Red dots represent significantly upregulated and blue dots represent significantly downregulated DEGs with p-value >4. **C-E.** qPCR analysis of C. PPARγ, D. CEBPα, and E. Fabp4 in, *in vitro* differentiated primary preadipocytes with adipogenic differentiation medium containing pyruvate concentrations ranging from 0.02 to 10 mM. **F-H.** qPCR analysis of F. PPARγ, G. CEBPα, and H. Fabp4 in, *in vitro* differentiated primary preadipocytes with adipogenic differentiation medium containing 4 mM pyruvate, analyzed at four-time points post-adipogenic induction. **I-K.** I. Oil Red O staining of lipid droplets (stained red) in ASCs during adipogenic differentiation with 4 mM of pyruvate at indicated time points was visualized by microscopy. J. Oil O Red destaining/elution absorbance was measured at 510 nm and normalized to the undifferentiated cells. K. Quantification of intracellular TG content. **L-N.** L. Oil Red O staining of lipid droplets (stained red) in bmMSCs during adipogenic differentiation with 4 mM of pyruvate at indicated time points was visualized by microscopy. M. Oil O Red destaining/elution absorbance was measured at 510 nm and normalized to the undifferentiated cells. N. Quantification of intracellular TG content. Data are mean ± SEM; * p < 0.05, ** p < 0.01, *** p < 0.001. Scale bar, 100 µm. (n = 3).

We next sought to employ differentiated murine bone marrow mesenchymal stem cells (bmMSCs) and adipose-derived stem cells (ASCs) to evaluate pyruvate’s role in intracellular lipid accumulation and triglycerides (TG) accumulation. They were investigated by Oil Red O (ORO) staining and colorimetry respectively. Both ASCs and bmMSCs were subjected to adipogenic induction with or without pyruvate for 8 days and 14 days respectively to determine which stage was most influenced by pyruvate. The ASCs staining revealed that pyruvate treatment reduced lipid vacuoles at each time point (**Fig. 1I**). The quantification of ORO stain also validated the significant decline in elution of the pyruvate-treated groups at each time point (**Fig. 1J**). This observed TG accumulation was corroborated by TG quantification, which demonstrated markedly reduced TG content with pyruvate (**Fig. 1K**). Similarly, bmMSCs results substantiated these effects of pyruvate with significantly decreased intracellular lipid accumulation (**Fig. 1L**), reduced ORO elution (**Fig. 1M**), and TG accumulation (**Fig. 1N**). These results indicate that pyruvate substantially inhibited adipogenic differentiation regardless of cell type.

The pathway analysis using bulk RNA sequence also verified that adipogenesis-related pathways viz., PPARg signaling, adipogenesis, cytokine & inflammatory response were significantly downregulated by pyruvate when the analysis was compared between the KEGG pathway and Wikipathway (**Fig. S1B**). UMAP analysis was used to cross-validate the results. It sorted the similar gene sets and positioned them into a cluster where the darker and larger the point corresponded to the significant enrichment of that pathway. Accordingly, we found that there was substantial negative regulation reaching a significant value for all the pathways related to adipogenesis involving fatty acid biosynthesis, cytokines, inflammatory response, and differentiation of white and brown adipocytes as a response to pyruvate (**Fig. S1C**). The volcano plot provided an additional representation of the enrichment results showing that pyruvate negatively impacts PPARg signaling. In summary, the pathway analysis using the RNA-seq data indicated the role of pyruvate in inhibiting the adipogenesis process (**Fig. S1D**).

### Pyruvate prevents HFD-induced weight gain and regulates white adipose tissue inflammation

To investigate the effect of pyruvate on the development of obesity *in vivo*, male C57BL/6 mice were fed on a chow diet (CD), High-fat diet (HFD), or HFD +Pyr or HFD + Orlistat for 12 weeks. Various morphological, physiological, and serological indicators were assessed. Starting from week 1, both HFD and CD mice increased body weight over time. The body weight of HFD-fed mice was significantly higher than their chow-fed counterparts at week 10 (**Fig. 2A**). This difference in weight gain was significant starting from the 4^th^ week of HFD supplementation between different groups, which was apparent even by the naked eye. Oral administration of pyruvate significantly suppressed HFD-induced body weight gain. The Orlistat group was used as the positive control. Besides body weight, another morphological characteristic that was evaluated was the abdominal circumference to gauge the central obesity of HFD-induced mice. As anticipated, the abdominal circumference of HFD-induced mice was significantly higher compared to the CD group. However, the administration of pyruvate significantly reduced central adiposity. A similar trend was also seen with orlistat, although it was more significant than pyruvate (**Fig. 2B**). We further analyzed the WAT depot to establish a source for enhanced weight gain in HFD mice as fat accumulation was the most dramatic change observed. Also, the distribution of WAT into visceral (vWAT) and subcutaneous (sWAT) depots are directly linked to metabolic diseases in obesity [50]. As observed, the mice fed with HFD showed a significant increase in vWAT which correlated with an increased body weight and central adiposity. This vWAT expansion displayed a reduced mass in the pyruvate-administered group. Interestingly, the sWAT depot mass was also increased in weight in the HFD group compared to the CD counterparts, but this expansion was significantly less compared to vWAT in male mice. Hence, the overall WAT depot mass correlated with adipocyte formation in the HFD group, which was significantly reduced by pyruvate administration (**Fig. 2C**). The brown adipose tissue did not show any significant weight difference (data not shown). Nevertheless, throughout the experimental feeding, the food intake between different groups was fundamentally the same (**Fig. 2D**).

**Figure 2:**
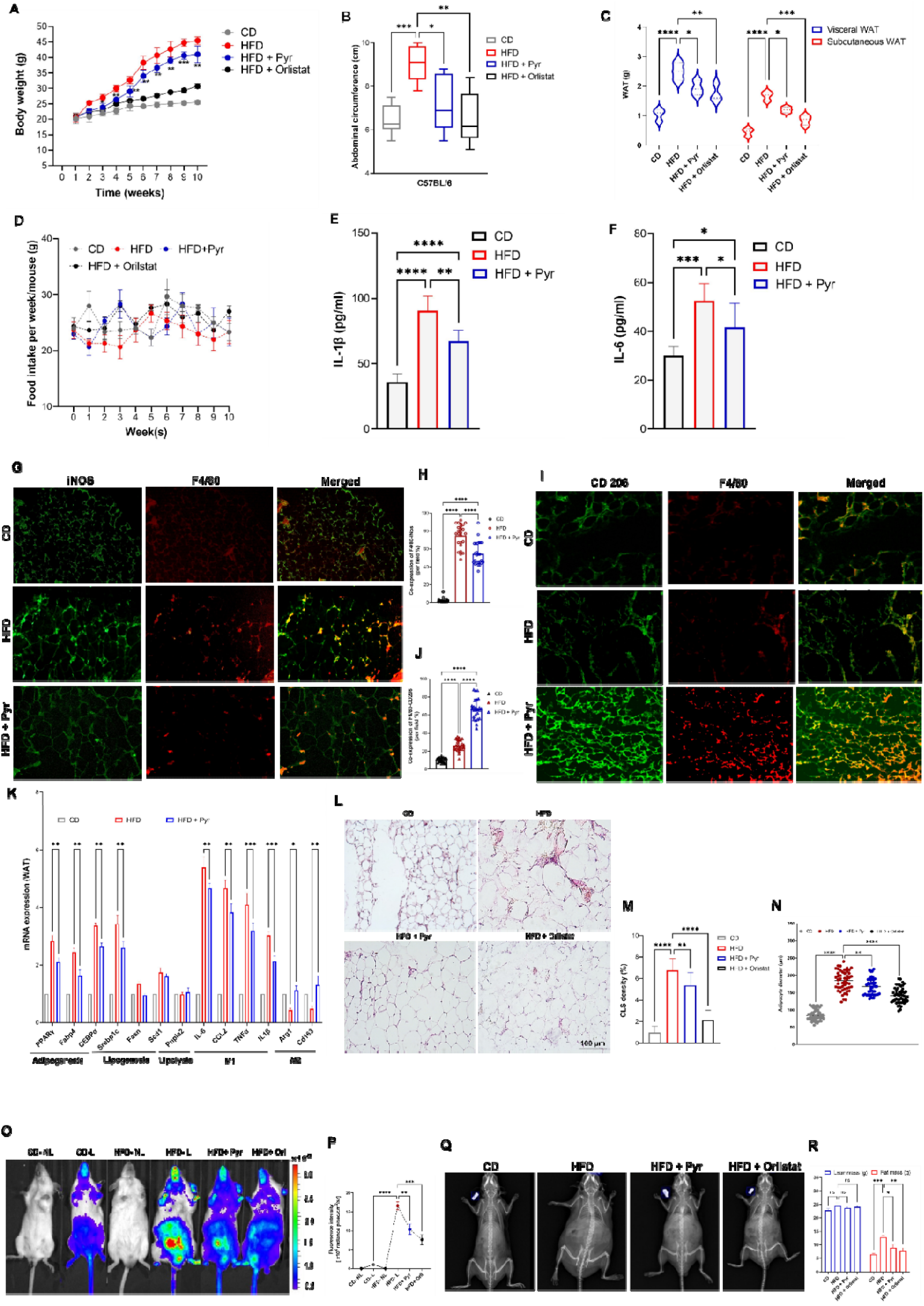
Pyruvate mitigates HFD-induced weight gain and white adipose tissue inflammation. C57BL/6 male mice were fed a high-fat diet (HFD) and chow diet (CD) for 10 weeks ± intervention with 1% pyruvate in drinking water. The experimental groups are mice fed on chow diet (CD), High-fat diet (HFD), HFD receiving pyruvate (HFD + Pyr), and HFD receiving orlistat (HFD + Orlistat). **A.** Weight gain in male mice. **B.** Abdominal circumference at the end of the experimental period. **C.** Comparison between adipose tissue (vWAT and sWAT) weight. **D.** Average food consumption per week. **E-F.** Serum level of inflammatory cytokines; E. IL-1β and F. IL-6 in indicated experimental groups. **G-J.** Representative immunofluorescence microphotographs of vWAT sections from three indicated groups; double stained with G. F4/80 (red) and iNOS (green) (M1 macrophage) H. Quantification of co-expression per field for M1 or I. F4/80 (red) and CD206 (green) (M2 macrophage) J. Quantification of co-expression per field for M2 macrophages. **K**. Quantitative RT-PCR of various genes in vWAT in indicated experimental groups, with the expression of CD-fed mice, normalized against *Gapdh*, being regarded as 1. **L-N**. L. Representative microphotographs of hematoxylin-eosin stained vWAT sections from different experimental groups. M. Histological quantification for Cown-like structures (CLS) and N. Comparison of mean adipocyte diameter, using Image J software. **O-P**. *In vivo* imaging of anesthetized NF-κB-Luc mice fed on HFD and CD for 10 weeks with/without 1% pyruvate in drinking water after 10 weeks. O. The figure shows representative bioluminescence images of chow diet-fed mice with no luciferin injection (CD-NL), chow diet-fed mice with luciferin injection (CD-L), high diet-fed mice with no luciferin injection (HFD-NL), high diet-fed mice with luciferin injection (HFD-L), high diet-fed mice receiving pyruvate with luciferin injection (HFD + Pyr) and high diet-fed mice receiving orlistat with luciferin injection (HFD + Pyr). The heat map depicts a two-dimensional visualization of light emitted ventrally from the mice following luciferin injection. All images were captured 10 minutes post-substrate administration with a 60-second exposure. Grayscale images were obtained prior to luminescence imaging as a reference. P. Quantification of *in vivo* NF-κB activity by measuring bioluminescence from NF-κB-Luc. **Q-R.** Q. Dual-energy-X-ray absorptiometry (DXA) scan images for indicated experimental groups. R. Lean mass and fat mass measurements by DXA for different experimental groups. Data are mean ± SEM; * p < 0.05, ** p < 0.01, *** p < 0.001. (n=6). Scale bar, 100 μm.

Obesity is associated with several comorbidities and low-grade chronic inflammation is one of them [51]. Hence, we next evaluated the plasma level of proinflammatory cytokines such as IL-1β and IL-6. We observed that in HFD-fed mice the secretion of these cytokines was substantially enhanced which was resolved in the pyruvate-administered group because the secretory levels of these cytokines were significantly reduced (**Fig. 2E-F**).

WAT is the primary location of inflammatory response in obesity with progressive immune cell infiltration [52]. Hence, we next conducted immunofluorescence analysis to examine polarized phenotypes of adipose tissue macrophages (ATMs) in vWAT as they are the most characteristic immune cells [16, 53]. F4/80 is a unique marker of murine resident tissue macrophages. Consequently, F4/80^+^iNOS^+^ double positive stain (merged) represented classically activated M1 macrophages, while F4/80^+^CD206^+^ double positive stain (merged) represented alternatively activated M2 macrophages. The immunofluorescence double staining uncovered some noteworthy changes in macrophage subtypes. We observed that M1 macrophages dramatically increased in the obese mice as compared to the lean mice and were significantly reduced in the pyruvate-administered group (**Fig. 2G-H**). However, M2 macrophages displayed an obvious decrease in obese mice but a dramatic increase in pyruvate treated group (**Fig. 2I-J**), this trend was consistent with previous studies [54]. Suggesting that diet-induced obesity leads to a shift in the activation state of ATMs from an M2-polarized state in lean animals to an M1 proinflammatory state in obese mice. However, pyruvate treatment can induce a switch back from M1-M2 recuperating towards a leaner state [54, 55].

To further confirm the effect of pyruvate on WAT inflammation, we assessed the expression level of M1 and M2 macrophage-related cytokines along with the marker genes for adipogenesis, lipogenesis, and lipolysis. qPCR analysis of vWAT revealed that the mRNA expression of M1 macrophage-related pro-inflammatory markers (IL-6, Ccl2, TNFα, and IL-1Β) was consistently elevated in HFD-induced obese mice as compared to the CD-fed lean mice but not the M2 macrophage related anti-inflammatory markers (Arg1 and Cd163). In fact, M2 markers showed a reverse trend where they were reduced in the HFD-fed obese mice. However, pyruvate treatment drastically diminished the expression of M1-related proinflammatory markers and increased the M2-related anti-inflammatory markers in the HFD group correlating to the previous results of macrophage polarization. In addition, the expression level of adipogenic markers (Pparg, Fabp4, and CEBPα) was found to be increased in the HFD-induced obese mice corresponding to enhanced adipogenesis. However, in the pyruvate-administrated group, mRNA expression of these markers was evidently reduced, relating to its anti-adipogenic effects. Interestingly, the lipogenic (Srebp1c, Fasn, and Scd1) as well as lipolytic (Pnpla2) marker expression level was rather unchanged in either of the groups as compared to the lean mice (**Fig. 2K**).

Other hallmarks of adipose-tissue inflammation are the greater incidence of crownLJlike structures (CLS) in WAT and adipocyte hypertrophy (increased size of adipocytes due to WAT expansion). CLS is characterized by the clustering of macrophages around a dying (or dead) adipocytes [56]. Histological analysis of vWAT exhibited a higher presence of CLS in WAT from that the HFD-fed obese mice group had remarkably high CLS as compared to the vWAT from CD-fed lean mice. (**Fig. 2 L-M**). Additionally, mice on HFD also showed a significant increase in the adipocyte size in vWAT when compared to the CD group (**Fig. 2N**). However, pyruvate administration substantially reduced both CLS as well as adipocyte diameter (hypertrophy) significantly as compared to the HFD group.

After assessing the low-grade systemic inflammation and the white adipose tissue inflammation by infiltrated macrophages in an obese state, the next plausible evaluation was for nuclear factor kappa B (NF-κB) activation and signaling NF-κB is crucially implicated in inflammatory processes accountable for the development of obesity [57], and its signaling activity is known to be elevated in mice fed HFD [58, 59]. Hence, we investigated the progression of NF-κB activity in mice fed on HFD and CD for 10 weeks using NF-κB luciferase reporter by employing IVIS (**Fig. 2O-P**). The *in vivo* imaging showed that mice without luciferase injection (CD-NL and HFD-NL) did not give any bioluminescence signal. While the signal from CD-fed mice with luciferase injection (CD-L) was considered the baseline signal (**Fig. 2O**). After taking out the baseline signal from other groups, we found that the overall NF-κB activity was elevated in HFD-fed mice. Interestingly, this signal was more intense around the abdomen, possibly derived from inflamed visceral WAT depots from the central adiposity. Nevertheless, mice on HFD that were orally administered with pyruvate showed a diminished bioluminescence signal relating to reduced NF-κB activity as compared to the HFD group. The orlistat group (positive control) showed expected results with a weakened NF-κB activity signal in the reporter mice (**Fig. 2P**).

Finally, to conclude that the decrease in body weight by pyruvate was due to a reduced accumulation of fat, we assessed whole-body fat mass (lean mass and fat mass) by performing dual-energy X-ray absorptiometry (DXA) for small animals (**Fig. 2Q**). The mice from the HFD group had a significantly higher fat mass at the end of the 10-week intervention as compared to mice fed with CD. In contrast, the orally administered pyruvate group fed on HFD resulted in a significantly ameliorated fat mass at the same 10-week time point (**Fig. 2R**). DEXA imaging confirmed that the increase in body fat occurred predominantly in the abdominal area, verifying visceral obesity. Hence, these results indicated that pyruvate-mediated attenuation in the HFD-induced weight gain could be attributed to its ability to decrease adipose tissue weight, regulate macrophage polarization, diminish NF-κB activity and resolve the overall inflammatory state of WAT.

As activation of the NF-κB transcription factor is linked to nuclear translocation of the p65 component, we also assessed the effect of pyruvate on p65 nuclear translocation *in vitro*. For this, TNFα was used to stimulate RAW macrophage cells, which resulted in a robust p65 nuclear translocation, which is otherwise confined in the cytoplasm under unstimulated conditions. In contrast, pyruvate-treated cells suppressed the nuclear translocation of p65 and most of the p65 were confined in the cytoplasm. (**Fig. S2A).** The quantification of confocal imaging demonstrated no overlap between green (cytoplasmic) and blue (nuclear) lines in the unstimulated group indicating no translocation. While, in the TNFα –stimulated group, the two lines predominantly overlapped marking nuclear translocation of the cytoplasmic fraction. However, in the pyruvate-treated group, the green (cytoplasmic) and blue (nuclear) lines were partially overlapped suggesting suppression of nuclear translocation as compared to the stimulated group (**Fig. S2B**). We further confirmed these results using a cell fractionation assay by extracting cytoplasmic and nuclear fractions at indicated time points. Western blot analysis revealed considerably less as well as postponed p65 translocation to the nucleus after TNFα stimulation in pyruvate-treated cells compared with untreated stimulated cells (**Fig. S2C-E**).

To further validate pyruvate’s role in suppressing inflammation by inhibiting NF-kB signaling, we used different cell types and aimed to evaluate the induction of IL-1β and IL-6, as it is essentially dependent on NF-κB [58]. Initially, we used RAW macrophage cells stimulated with TNFα. The qPCR and ELISA demonstrated that both low (2mM) and high (4mM) concentrations of pyruvate inhibited TNFα-induced mRNA expression levels (**Fig. S2F-G**) as well as the secretory level (**Fig. S2H-I**) of IL-1Β and IL-6 in a dose-dependent manner respectively. We next utilized *in vitro* differentiated BMMLJ from TNF-tg mice to mimic the systemic inflammatory state. The qPCR revealed a comparable dose-dependent reduction in the mRNA expression of IL-1β and IL-6 (**Fig. S2J-K**). In addition to mRNA level, we also measured secreted cytokines expression from macrophages stimulated by TNFα via ELISA and observed that IL-1β and IL-6 expression in the supernatants was also significantly reduced by pyruvate in a dose-dependent manner (**Fig. S2L-M**). Finally, we used *in vitro* differentiated primary BMMLJ from C57BL/6 mice stimulated with TNFα. Both qRT-PCR and ELISA demonstrated that pyruvate dose-dependently ameliorated TNFα-induced cytokine mRNA expression(**Fig. S2N-O**) and release (**Fig. S2P-Q**) of IL-1β and IL-6, respectively.

Macrophage polarization is strongly associated with inflammation where M1-type macrophages play a pivotal role in the pro-inflammatory, and M2-type macrophages contribute to anti-inflammatory responses. The M1*/*M2 polarization profile of macrophages with or without pyruvate was investigated using *in vitro* differentiated primary BMMLJ. We measured the expression of cytokines in different macrophage phenotypes. The mRNA expression of M1-related cytokines IL-6 and iNOS was dramatically attenuated by pyruvate significantly (**Fig. S2R**) whereas the expression of M2-related cytokines Arg1 and Mgl1 were enhanced by pyruvate (**Fig. S2S**). These results indicated that pyruvate induces a phenotypic switch from M1(pro-inflammatory) to M2 (anti-inflammatory) phenotype resolving the overall state of inflammation.

### Pyruvate ameliorates HFD-induced metabolic dysregulation and hepatic steatosis

To test whether HFD impaired glucose tolerance, an oral glucose tolerance test (OGTT) was performed. Followed by an overnight fast, glucose (mg/dL) levels were quantified in a fasting state and 15, 30, 45, and 60 min after administering glucose solution via oral gavage (**Fig. 3A**). The fasting glucose levels remained unchanged between experimental groups (**Fig. 3B**). However, after the glucose challenge, the blood glucose concentration reached its peak after 15 min in all groups. The HFD group showed a persistently elevated glucose concentration compared to that of the CD group followed by a delayed decrease, demonstrating a slow glucose clearance from the bloodstream. These findings suggested that HFD feeding for 10 weeks was able to induce glucose intolerance in mice. The HFD-fed mice treated with pyruvate showed a sharp decline in blood glucose levels after glucose load at 30 min and continued to reduce these levels significantly as compared to HFD-fed mice. The orlistat group also showed the same trend where 60 min time point showed glucose levels comparable to the CD-fed mice. Regulation of glucose and lipid metabolism are intricately interconnected [36]. Hence, to investigate whether HFD could induce dyslipidemia, we examined the serum lipid profile in HFD-fed mice and other experimental groups, at the end of the experimental period. Serum TG, TC, LDL, and HDL levels were measured and found to be significantly higher in the HFD group as compared to the CD group (**Fig. 3C-E**). Nevertheless, oral administration of pyruvate showed anti-hyperlipidemic activity by decreasing TG, TC, and LDL levels with significant differences. The level of HDL was found to be decreased in HFD-fed mice, and remained unchanged by pyruvate however, orlistat resulted in the significant elevation of HDL (**Fig. 3F**) and reduction of other lipid markers.

**Figure 3:**
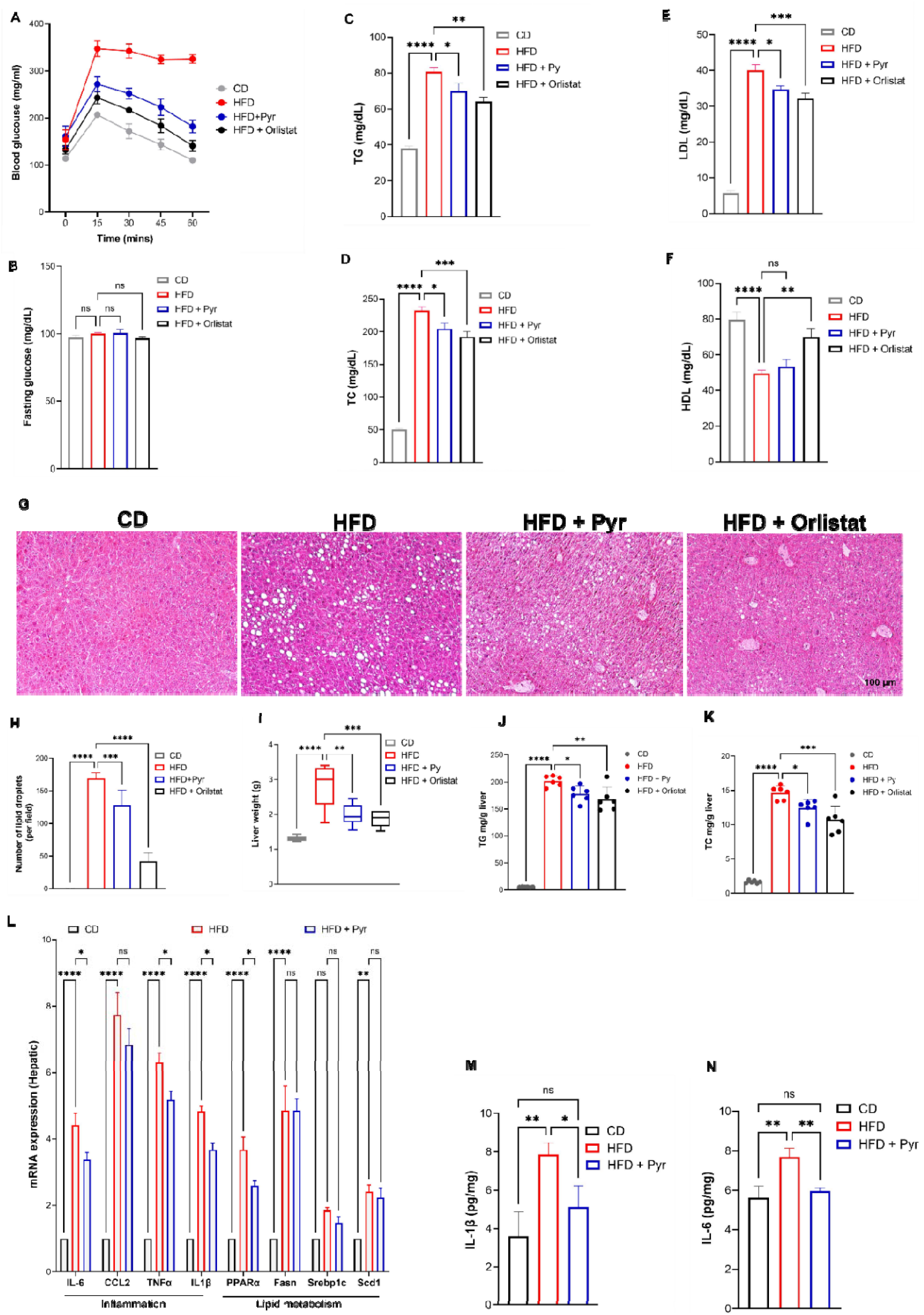
Pyruvate prevented HFD-induced metabolic dysregulation and hepatic steatosis. **A.** Blood glucose level measured by OGTT after overnight fasting in the indicated experimental groups. **B.** Fasting blood glucose level comparing different experimental groups. **C-F**. Serum lipid parameters of indicated experimental group C. Triglycerides level (TGs) D. Total Cholesterol (TC) E. Low-density lipoprotein cholesterol (LDL) F. High-density lipoprotein cholesterol (HDL). **G-H.** G. Hematoxylin and eosin-stained liver sections representing mice from each experimental group. H. Quantification of the lipid droplets (steatosis) for each group. **I.** Weight of liver tissue. **J.** Hepatic triglyceride level. **K.** Hepatic total cholesterol level. **L**. Quantitative RT-PCR of the mRNA levels of inflammation and lipid metabolism markers in the liver, as indicated. The expression of CD-fed mice, normalized against *Gapdh*, is regarded as 1. **M-N**. Hepatic mRNA expression of inflammatory genes M. IL-1β and N. IL-6. Data are mean ± SEM; * p < 0.05, ** p < 0.01, *** p < 0.001. (n=6). Scale bar, 100 μm.

Metabolic disorders including obesity, inflammation, hyperglycemia, and dyslipidemia, are usually comorbid with nonalcoholic fatty liver disease (NAFLD) [37]. NAFLD is characterized by intrahepatic triglyceride fat accumulation in the liver also termed hepatic steatosis [38]. The histological analysis was performed to investigate the role of pyruvate on hepatic lipid accumulation (Fig. 3G). Hematoxylin-eosin (H&E) staining of livers in the HFD group displayed remarkable vacuolation indicative of lipid deposition inside the liver parenchyma (**Fig. 3G**) as compared to the CD-fed mice. Nevertheless, the degree of HFD-induced hepatic steatosis was significantly attenuated in pyruvate– and orlistat-administered mice as evidenced by reduced lipid vacuoles in the liver(**Fig. 3H**).

Consistent with the results of histological analysis, the liver weight in pyruvate-administered mice was substantially lower than that in high-fat diet-fed mice suggestive of lipid accumulation (**Fig. 3I**). To further investigate the therapeutic role of pyruvate on hepatic lipid profiles, we conducted a biochemical analysis. Compared with CD-fed mice, the HFD group exhibited elevated hepatic TG and TC contents (**Fig. 3J-K**). However, pyruvate administration significantly improved hepatic lipid profiles under HFD conditions. We next examined the expressions of genes associated with proinflammatory cytokines including as well as in lipid metabolism by RT-PCR analysis. We observed that the expressions of proinflammatory cytokines including IL-6, CCL2, TNF-a, and IL-1β were increased in the HFD group compared to the CD-fed mice. Whereas pyruvate administration inhibited HFD-induced hepatic inflammatory responses by significantly downregulating the expression of proinflammatory genes except for CCL2. Additionally, the expressions of lipid metabolism-associated genes including PPARα, Fasn, Scd1, and srebp1c, were remarkably higher in the HFD group as compared to the CD-fed group. Interestingly, there was a non-significant downregulation in their expression after pyruvate administration except PPARα, which showed significantly attenuated expression levels. (**Fig. 3L**). Lastly, the levels of IL-1β and IL-6 were evaluated in the livers using ELISA. The results indicated that the levels of these proinflammatory cytokines were higher in the HFD-induced obese mice compared to the normal CD-fed control group. However, pyruvate significantly reduced their levels in the liver (**Fig. 3M-N**). Taken together, these results indicated that pyruvate prevented the pathological process of NAFLD by decreasing lipid deposition and inflammatory responses and maintained the overall hemostasis without directly affecting lipid metabolism.

Gender-based differences in adipose tissue distribution have been well documented. Similarly, sex differences can lead to disparity in drug responses [39]. Thus, to determine if pyruvate also attenuated HFD-induced pathologies in – C57BL6/J female mice, we fed the mice on HFD or CD for 10 weeks. Female mice on HFD showed an increase in weight during the studied period of 10 weeks, which was significantly higher than mice on the control mice fed on the CD diet (**Fig. S3A**). However, the overall increase in body weight was accentuated in male mice on the HFD diet. Pyruvate-administered mice showed a considerable decline in weight gain which was significantly lower than HFD-fed mice. Consistent with the body weight, the abdominal circumference of female mice on HFD was significantly higher as compared to the CD-fed mice. However, the pyruvate-administered mice showed significantly less circumference as compared to the HFD group (**Fig. S3B**). However, the overall increase in the abdominal circumference was more in male mice as compared to female mice.

Following, we measured the fat pad weight of the vWAT and the sWAT depots in all experimental groups, and found that upon HFD feeding, females accumulated comparable vWAT and sWAT depots, unlike male mice where vWAT was considerably higher than sWAT in both CD– and HFD-fed conditions (**Fig. S3C**).

In OGTT, plasma glucose concentrations showed a similar trend as in male mice where the peak was attained after 15 minutes of a glucose load, which was suggestively higher in HFD mice than in CD-supplemented mice and remained elevated at a significantly higher level than CD-supplemented mice, demonstrating impaired glucose tolerance. While in pyruvate-administered mice, a gradual decline was observed after striking the peak and at 60 min there was a non-significant difference compared to the control CD mice. Nevertheless, this impairment was overall enhanced in male mice on HFD (**Fig. S3D**). However, the fasting glucose levels in female mice remained constant throughout the groups (**Fig. S3E**). In terms of dietary intake, no disparities were observed between all experimental groups (**Fig. S3F**). Circulating levels of serum TG, TC, and LDL levels were measured and found to be significantly high in response to the HFD (**Fig. S3G-I**), and HDL was decreased as compared to the CD group (**Fig. S3J**). This trend was the same as in male mice, but they displayed a heightened elevation of circulatory lipids. By attenuating the concentration of circulating lipids, pyruvate also had the same results in female mice, with no significant effect on HDL.

H&E staining revealed a significant enhancement in vacuolation in response to HFD in female mice over their respective CD-fed controls. However, in liver sections from pyruvate-treated mice, lipid deposition was reduced substantially. Moreover, the hepatic changes were comparable between male and female HFD-supplemented mice. (**Fig. SK-L**). Reverberating with histological analysis, liver weight in HFD mice was higher than CD control mice, but mice receiving pyruvate exhibited lower liver weight in HFD fed state (**Fig. S3M).** Finally, it was noticed that female HFD-fed mice exhibited hepatic hyperlipidemia while pyruvate reduced their elevated levels in the HFD state, based on the biochemical analysis of hepatic TG and TC **(Fig. S3N-O**). Orlistat showed large-scale therapeutic effects on the analysis of HFD-induced changes and pathologies. Hence, these results clearly indicate that despite the mice being sexually dimorphic with respect to fat mass distribution in obesity, the pyruvate effect in neutralizing the pathophysiology is preserved.

By evaluating various physiological parameters in non-obese CD-fed mice with and without pyruvate, we excluded the possibility of adverse effects of pyruvate on healthy mice. The non-obese CD-fed mice administered with or without pyruvate did not show any difference in body weight (**Fig. S4A**) and other parameters like abdominal circumference (**Fig. S4B**). The oral glucose test also demonstrated the same trend of glucose clearance from the bloodstream in both groups (**Fig. S4C**). The H&E staining of WAT showed comparable adipocyte diameter (**Fig. S4D-E**) along with comparable fat pad weights of vWAT and sWAT (**Fig. S4F**). The liver and kidneys are common organs affected by chemical toxicity. Hence, it was imperative to evaluate the effect of pyruvate administration on vital organs (**Fig. S4G**). Undoubtedly, the systemic administration of pyruvate presented no toxicity in vital organs where the normal anatomy of each indicated organ was retained. In the liver, hepatocyte size was analogous, and no derangement of liver enzymes was found (data not shown). A comparable white-to-red pulp ratio was seen in the spleen, as well as a healthy glomerular area and podocytes in the kidney. The heart also displayed normal anatomical features with typical cardiomyocyte area comparable between both groups (**Fig. S4G**). Hence, pyruvate did not exhibit any undesirable effects, indicating its safety as a dietary supplementation and/or drug candidate. Consistent with the histological analysis, there was no difference in the weight of these organs between the two experimental groups (**Fig. S4H).** Food intake demonstrated marked daily variation; however, the overall difference was nonsignificant (**Fig. S4I**). Finally, the mice consumption of water was comparable with or without pyruvate in the water, indicating an absence of aversion to pyruvate-containing water (**Fig. S4J**).

### cPLA2 is a previously unrecognized direct target of pyruvate

During our efforts to understand how pyruvate inhibits obesity and inflammation at the molecular level, we focused on identifying its direct binding targets. For this purpose, a drug affinity responsive target stability (DARTS) assay was used with its strategy outlined (**Fig. 4A**). The cell lysate was treated with or without pyruvate and this protein mixture was separated by sodium dodecyl sulfate-polyacrylamide gel electrophoresis (SDS-PAGE) after proteolytic digestion of the cell lysates. The results demonstrated that pyruvate protected a band with a molecular weight of approximately 80 kDa **(Fig. 4B).**

**Figure 4:**
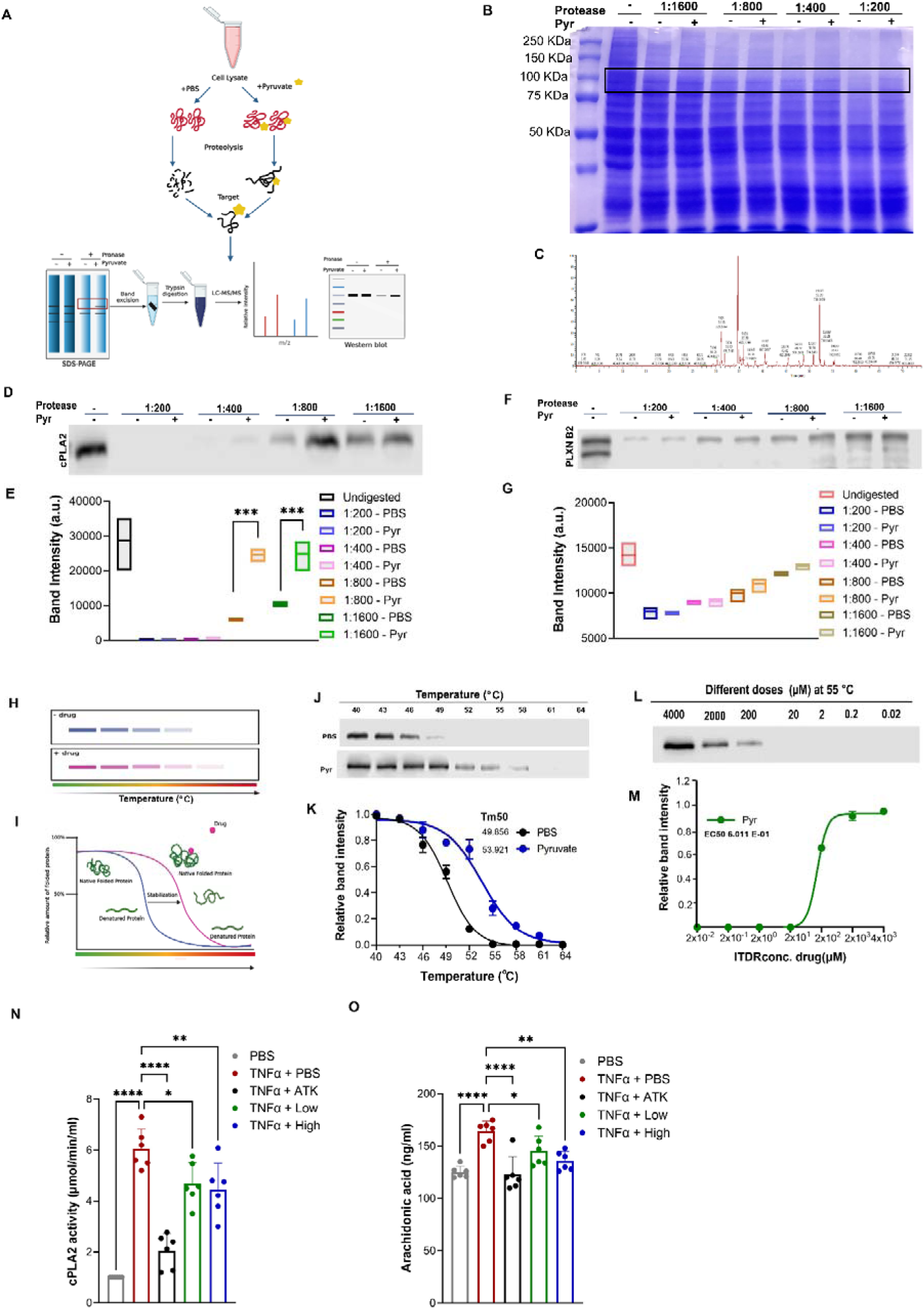
Target identification and validation for the molecular target of pyruvate. **A.** Schematic representation of drug affinity responsive target stability (DARTS). **B.** Coomassie blue staining of DARTS assay gel. The band with a molecular weight of approximately 80 kDa was protected by pyruvate treatment (indicated in a black box). **C.** Mass spectrometry adapted image for *PLA2G4A*, encoding cPLA2. **D-E.** D. DARTS followed by immunoblot employing anti-cPLA2 antibody to confirm pyruvate’s binding target. E. Quantification of panel D showing relative cPLA2 band intensity in with or without pyruvate groups. **F-G.** F. DARTS followed by immunoblot employing anti-PLEXINB2 antibody to confirm pyruvate’s binding target. G. Quantification of panel F showing relative PLEXINB2 band intensity in with or without pyruvate groups. **H**. Schematic representation of CETSA. **I.** Schematic representation associated with CETSA outlining the principle. **J-K.** J. CETSA melt response at indicated temperatures with or without pyruvate using western blot analysis. K. CETSA associated densitometry analysis curve. **L-M.** L. Isothermal dose response (ITDR) with serial concentrations of pyruvate at 55°C. M. ITDR associated curve showing relative soluble fraction using western blot analysis. **N.** cPLA2 activity of RAW264.7 cells transfected with cPLA2 expression plasmid, treated with TNFα, ATK, or pyruvate overnight. **O**. Arachidonic acid (AA) levels: BMDMs without or with TNFα in the absence or presence of pyruvate for 48 hours were examined for AA levels using an ELISA kit. ATK was used as a positive control. Data are mean ± SEM; * p < 0.05, ** p < 0.01, *** p < 0.001. (n=3).

We excised this band and subjected it to mass spectrometry for identification of the protein (**Fig. 4C**). Consequently, PLA2G4A encoding cPLA2 and PLXNB2 encoding for Plexin B2 protein were identified as potential targets. As a means of confirmation of these targets, western blot analysis of DARTS samples with different protease-to-cell lysate ratios revealed that pyruvate protected the cPLA2 band (**Fig. 4D-E**) but not the PLEXNB2 band (**Fig. 4F-G**) against enzymatic digestion.

As a means of further validating the association between pyruvate and cPLA2, CETSA was performed. With CETSA, it is possible to quantify the change in thermal denaturation of a target protein under varying changes in temperature and concentration of the compound of interest (**Fig. 4H-I**). Pyruvate prevented the denaturation of cPLA2 and maintained a higher level of cPLA2 in the soluble state under several temperatures, most evident at 49°C when compared with PBS **(Fig. 4J).** As shown by the melt curve, pyruvate caused a significant shift and change in Tm (Tm = 53.921°C) in comparison with PBS (Tm = 49.85°C) (**Fig. 4K**).

We evaluated CETSA’s performance at 55°C by analyzing its isothermal dose response (ITDR). According to these results, pyruvate protected cPLA2 denaturation in a dose-dependent manner, with an EC50 of 6.011E-01 (**Fig. 4L-M**). Taken together, cPLA2 has been identified as a target of pyruvate that has previously not been recognized.

Following confirmation and validation of our target, we next evaluated the effects of pyruvate on cPLA2 activity. In this assay, we used arachidonyl trifluoromethyl ketone 27 (ATK), a known cPLA2 inhibitor as a positive control. Results indicated that pyruvate, similar to ATK, reduced the activity of cPLA2 induced by TNFα in a dose-dependent manner (**Fig. 4N**). It has been shown that inflammatory conditions, such as elevated TNFα, promote the translocation of cPLA2 to intracellular phospholipid membranes [19]. cPLA2 is primarily responsible to promote phospholipid hydrolysis to produce arachidonic acid (AA),which, in turn, is a known mediator of obesity and NF-κB signaling [40]. Accordingly, we examined whether pyruvate inhibited TNFα induced cPLA2-mediated AA production and the data revealed that pyruvate indeed reduced AA production significantly in a dose-dependent manner (**Fig. 4O**).

### cPLA2 is indispensable for pyruvate’s therapeutic effects in HFD-induced obesity

To determine the dependence of pyruvate’s effects on cPLA2 *in vivo*, we generated cPLA2 KO mice using a crossbreeding strategy (**Fig. S5A**), which was verified through genotyping (**Fig. S5B**). Furthermore, the global deletion of this gene was confirmed by measuring cPLA2 levels in various organs (spleen, kidney, heart, and liver) of KO mice (**Fig. S5C-D**). The age-matched mice (both WT and KO) were divided into different experimental groups according to their sexes after breeding and weaning cycles. In our further study design as illustrated (**Fig. S5E**), the control group of mice is fed a chow diet (CD) while the experimental group of mice is fed a high-fat diet (HFD) with or without pyruvate supplementation for a period of 16 weeks.

To examine the role of pyruvate in diet-induced obesity and its dependence on cPLA2 *in vivo*, we placed cPLA2 WT and cPLA2 KO mice on HFD for 16 weeks followed by the evaluation of various physiological and pathological processes. As compared to CD-fed mice, mice fed an HFD were significantly more likely to be obese and gain weight than mice fed a CD. By week six of the experiment, this difference in weight gain became apparent. Pyruvate supplementation significantly suppressed HFD-induced weight gain in WT mice, but strikingly, pyruvate was not effective in suppressing weight gain in cPLA2 KO mice (**Fig. 5A-B**). Further, it was noted that KO mice gained weight at a considerably slower rate than WT mice, with KO mice exhibiting a considerably slower tendency to gain weight.

**Figure 5:**
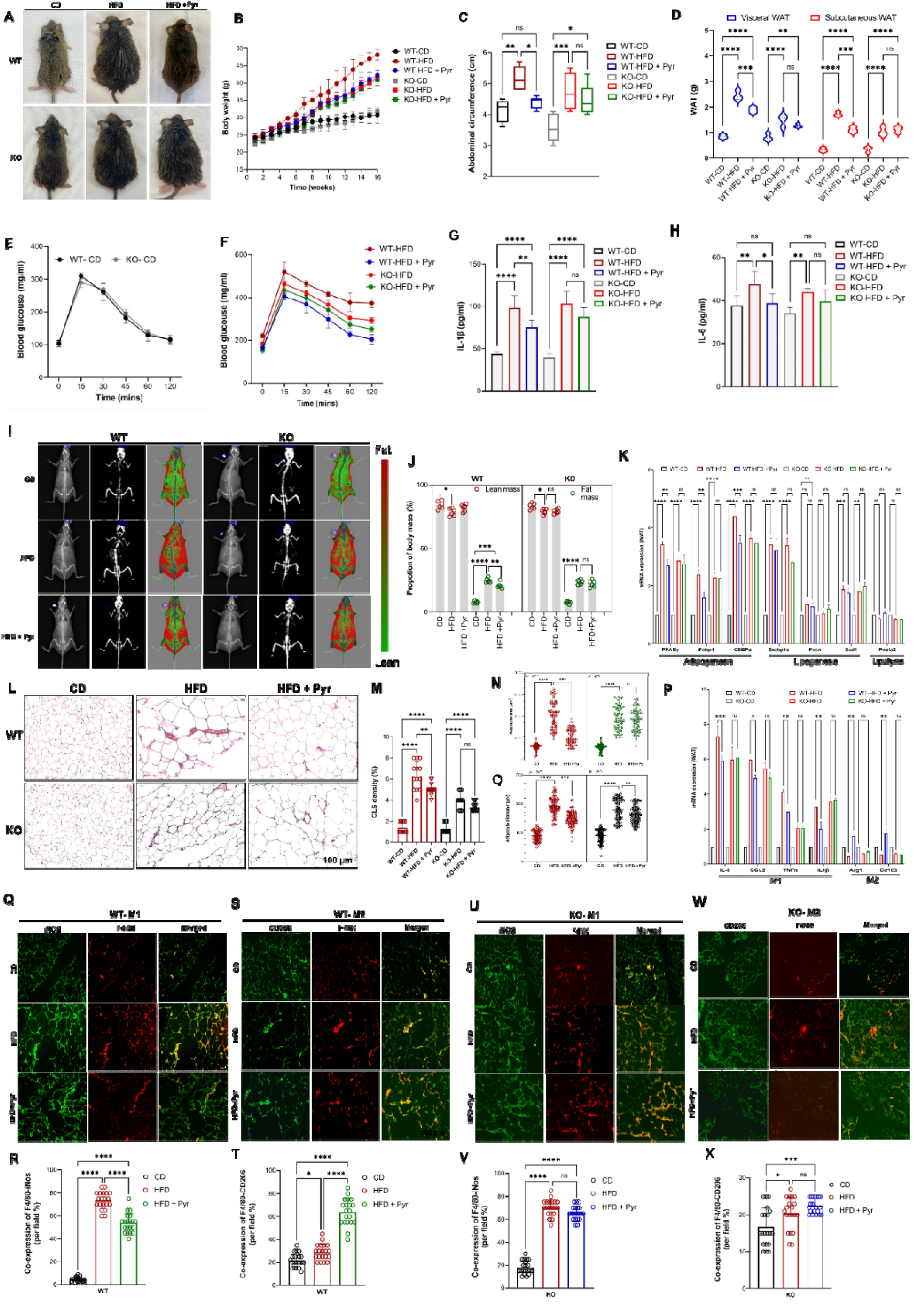
cPLA2 is essential for pyruvate’s anti-obesity effects. cPLA2 WT and cPLA2 KO mice were fed a CD or HFD for 16 weeks with or without pyruvate intervention. **A.** Gross appearance. **B.** Body weights for the indicated periods. **C**. Measurement of abdominal circumference. **D.** Comparison between adipose tissue (vWAT and sWAT) weight for the indicated groups. **E.** Blood glucose level measured by OGTT after overnight fasting in WT and KO mice fed on CD. **F.** Blood glucose level measured by OGTT after overnight fasting in the indicated HFD-fed experimental groups. **G-H.** Serum level of inflammatory cytokines; G. IL-1β and H. IL-6 in indicated experimental groups. **I-J.** I. Dual-energy-X-ray absorptiometry (DXA) scan images for indicated experimental groups showing (left to right) x-ray image, bone enhanced image, and color composition image (fat and lean mass indicated by red and green respectively). J. Body composition measurement by DXA showing lean mass and fat mass for different experimental groups. **K**. Quantitative RT-PCR of various target genes for adipogenesis, lipogenesis, and lipolysis in vWAT in indicated experimental groups on HFD with or without pyruvate, with the expression of CD-fed WT and KO mice, normalized with β*-actin*, being regarded as 1. **L-O.** L. Representative microphotographs of hematoxylin-eosin stained vWAT sections from different indicted experimental groups showing CLS. M. Histological quantification for CLS percentage N-O. Comparison of mean of N. adipocyte area and O. adipocyte diameter, using Image J software. **P.** Quantitative RT-PCR of various target genes for M1 (IL-6, Ccl2, TNFα, and IL-1β) and M2 macrophage markers (Arg1 and Cd206) in vWAT in indicated experimental groups on HFD with or without pyruvate, with the expression of CD-fed WT and KO mice, normalized with β*-actin*, being regarded as 1. **Q-T.** Representative immunofluorescence microphotographs of vWAT sections from cPLA2 WT mice from the indicated experimental group; double stained with Q. F4/80 (red) and iNOS (green) (M1 macrophage) R. Quantification of co-expression per field for M1 macrophages or S. F4/80 (red) and CD206 (green) (M2 macrophage) T. Quantification of co-expression per field for M2 macrophages. **U-X.** Representative immunofluorescence microphotographs of vWAT sections from cPLA2 KO mice from the indicated experimental group; double stained with U. F4/80 (red) and iNOS (green) (M1 macrophage) V. Quantification of co-expression per field for M1 macrophages or W. F4/80 (red) and CD206 (green) (M2 macrophage) X. Quantification of co-expression per field for M2 macrophages. Data are mean ± SEM; * p < 0.05, ** p < 0.01, *** p < 0.001. (n=6-8). Scale bar, 100 μm.

Often used as a marker of obesity in HFD animal models, abdominal circumference is measured in mice that have been fed a high-fat diet (HFD) to induce obesity. Consequently, when we compared this parameter between different groups, we found that the abdominal circumference of WT mice fed on HFD supplemented with pyruvate was significantly lower than that of the HFD group. However, in KO mice, pyruvate did not result in a statistically significant reduction in the girth of the abdomen (**Fig. 5C**).

Both vWAT and sWAT were significantly increased in HFD groups as compared to CD groups regardless of genotype. Although vWAT showed more increment than sWAT, both showed substantial attenuation following pyruvate treatment in WT. The effect of pyruvate on WAT depots in KO mice was abolished, and its supplementation was ineffective in reducing WAT deposits. Additionally, a significant difference was also observed between the WAT depots of the HFD and KO groups, indicating that cPLA2 plays a role in obesity (**Fig. 5D**). We found no significant weight difference in brown adipose tissue regardless of genotype or experimental group (data not shown).

From both WT and KO control groups fed on CD, the OGTT curve typically demonstrated a rapid increase in blood glucose levels shortly after oral glucose was administered. This was followed by a sharp decline in blood glucose levels indicating clearance of glucose from the bloodstream. This trend was strikingly similar between the two groups (**Fig. 5E**). In contrast, HFD mice from WT as well as KO showed an exaggerated initial peak at 15 min, which slowly started to decline around 30 minutes but failed to return to the baseline at the endpoint, indicating impaired glucose tolerance in HFD-induced obese mice. Although pyruvate administration decreased the elevated levels after 30 min and almost returned to baseline levels in WT, it failed to achieve the same trend in the cPLA2 KO group (**Fig. 5F**). The fasting glucose level was non-significantly increased in HFD-fed mice and was comparable between experimental groups (**Fig. S6C).**

To determine whether HFD could cause dyslipidemia in cPLA2 WT and KO mice, the serum lipid profiles in both experimental groups were examined. It was found that both the WT and KO groups of mice fed HFD had significantly elevated TG, TC, and LDL levels. In contrast, pyruvate administration resulted in decreased lipid levels in WT mice whereas pyruvate failed to restore the lower levels of these lipids in cPLA2 KO mice. (**Fig. S6D-F**). Despite this, HDL levels in WT were reduced after HFD, but in KO mice they were maintained at the same level as in control CD mice. The level of HDL was not increased by pyruvate regardless of genotype (**Fig. S6G**).

Obesity-associated chronic low-grade systemic inflammation is mainly mediated by WAT and plays an essential role in the pathogenesis of metabolic abnormalities [41]. Thus, we studied both systemic and adipose tissue inflammation. The plasma levels of proinflammatory markers IL-1β and IL-6 were increased in HFD mice as compared to CD controls in both WT and KO mice, suggesting systemic chronic low-grade inflammation. Significant attenuation of these inflammatory markers was observed in WT-HFD mice when pyruvate was administered. In contrast, pyruvate could not reduce these levels in the KO-HFD group (**Fig. 5G-H**).

We next performed dual-energy X-ray absorptiometry (DXA) to measure the *in vivo* body composition, including fat mass, and lean mass (muscle and bone) of mice in different experimental groups. In diet-induced obese mice, DXA-derived values were highly correlated with increased body weights, WAT content, and abdominal circumference after 16 weeks of HFD (**Fig. 5I-J**). As compared to control mice on a regular diet in both genotypes, HFD increased fat mass by more than 2 folds in both WT and KO mice, despite a modest difference in weight gain between the two genotypes. However, the lean mass demonstrated a slight decrease in HFD groups. The administration of pyruvate in cPLA2 WT mice exhibited consistent results with previous findings, including significant reductions in fat mass but did not alter the lean mass. In contrast, cPLA2 KO abrogated pyruvate’s anti-obesity effects and did not significantly reduce fat mass. In both WT and KO animals, pyruvate treatment did not affect lean mass and it remained marginally decreased. Surprisingly, a non-significant difference was found between high-fat diet mice and control diet (CD) mice in terms of bone area and bone mineral content (**Fig. S6 H-I**).

We next assessed the key adipocyte functions in vWAT from WT and KO experimental groups by measuring the expressions of markers for adipogenesis (Pparg and Fabp4 and CEBPα), lipogenesis (Srebp1c, Fasn, and Scd1), and lipolysis (Pnpla2). In both genotypes, HFD-receiving mice showed the highest expression levels of adipogenic markers; however, KO mice showed relatively lower levels than WT mice. Except for Fasn, other markers of lipogenesis (Srebp1c, and Scd1) were elevated, whereas the lipolysis marker was not significantly affected across all experimental groups. Adipogenic markers were robustly inhibited in the pyruvate-treated group in WT, while lipogenesis and lipolysis markers were unaffected, indicating that pyruvate primarily inhibited adipogenesis. As compared to WT, the vWAT from KO did not show any reduction in pyruvate-supplemented groups, including adipogenic markers, proving that cPLA2 plays a critical role in pyruvate’s mechanism of action. (**Fig. 5K**).

Our next consideration was that WAT browning contributes to a reduction in white adipose mass [42]. Since pyruvate administration reduced vWAT weight significantly, it was imperative to evaluate its effects on thermogenic markers. In both WT and KO mice, expressions of browning or thermogenic markers (UCP-1, Cidea, Elovl6, and Cox8b) were reduced following HFD feeding compared to CD-fed mice. Despite this, their expressions remained comparable across groups. It was noteworthy that neither WT nor KO group was affected by pyruvate in terms of suppressing or elevating these markers (**Fig. S6J**).

There is a common association between obesity and WAT inflammation, hence, we next performed a microscopic analysis of adipose tissue histology by H&E staining. There was no inflammatory cell infiltration within the adipose tissue of the CD group, which consisted of white adipocytes with a smooth thin layer of cytoplasm surrounding single spherical or oval-shaped lipid droplets. However, there was a significant increase in the size of individual adipocytes after 16 weeks of HFD, which was confirmed by measuring the diameter as well as the area of each adipocyte. Furthermore, CLS, a pathological hallmark of tissue inflammation indicating macrophage infiltration, was more prominent in the HFD group in both genotypes. Pyruvate demonstrated a promising anti-inflammatory role in WAT from the WT group, by reducing both the size of adipocytes as well as CLS. By knocking out cPLA2, this effect of pyruvate was abolished, and the area, diameter, and CLS density of the HFD group and the pyruvate-supplemented group were not significantly different (**Fig. 5L-O**).

We investigated the mRNA levels of M1 and M2 macrophages in vWAT of the WT and KO groups considering the increased CLS in WAT, which is a marker of macrophage infiltration. We observed that the expression levels of M1 macrophage markers (IL-6, Ccl2, TNFα, and IL-1β) were consistently elevated in vWAT of HFD-fed WT and KO mice compared to CD-fed control mice. However, the expression levels of M2 macrophage markers (Arg1 and Cd206) were decreased. In WT mice, pyruvate significantly reduced their expressions, but in KO mice, pyruvate failed to reach significance. Furthermore, anti-inflammatory M2 macrophage markers (Arg1 and Cd206) were dramatically reduced in HFD-fed mice as compared to the control. As expected, pyruvate reversed this modulation of gene expression in pyruvate-administered HFD-fed mice, where M2 levels were elevated significantly in WT mice, but not in cPLA2 ablated mice (**Fig. 5P**). These studies demonstrate that pyruvate has therapeutic potential for resolving obesity by favoring the polarization of M2 macrophages and inhibiting inflammation. These results also reiterate earlier findings that cPLA2 is fundamental for pyruvate’s physiological action.

Using immunofluorescence staining for the respective markers, we analyzed cells that were positive for classically activated M1 macrophages (F4/80^+^iNOS^+^) or alternatively activated M2 macrophages (F4/80^+^CD206^+^) to access macrophage infiltration in WAT of cPLA2 WT and KO mice from different experimental groups. Consistent with H&E observations, there were more proinflammatory macrophages in the WAT of WT mice after HFD feeding, as compared to CD-fed mice. A significant reduction in M1 macrophage levels was observed in WAT obtained from mice that had been administered pyruvate (**Fig. 5Q-R**). In contrast, the M2 anti-inflammatory macrophages were significantly increased in WAT isolated from HFD-fed pyruvate-administered mice which were otherwise extremely redundant in the WAT of obese mice (**Fig. 5S-T**). Thus, in the cPLA2 WT genotype, pyruvate administration reduced the development of chronic inflammation.

As observed by double-positive staining in WAT of KO mice, HFD promoted the M1 subtype and reduced the M2 subtype, indicating a shift toward a pro-inflammatory state compared with a CD-fed lean state. In contrast, the effects of pyruvate on regulating the M1 and M2 switch were completely disabled in KO mice, and M1 infiltration was comparable between KO-HFD and KO-HFD+Pyr, which is greatly increased as compared to the lean mice (**Fig. 5U-V**). In addition, KO mice showed a negligible number of M2 macrophages in the WAT isolated from both, HFD and HFD+Pyr (**Fig. 5W-X**). In summary, HFD disrupts the balance between M1 and M2 macrophages, promoting inflammation by favoring M1 macrophages regardless of genotype. Pyruvate restored this imbalance in WT mice convalescing towards a leaner state, but in CPLA2 KO mice this effect was abolished.

### cPLA2 ablation impairs pyruvate-mediated amelioration of hepatic lipid accumulation and dyslipidemia

As obesity is associated with a variety of alterations in hepatic parameters, we evaluated them including liver weight, inflammation, and fat accumulation (hepatic steatosis). As compared to CD-fed mice, liver weights in WT and KO mice on HFD were statistically higher. Nevertheless, the WT-HFD group showed a reduction in liver weight following pyruvate administration, suggesting reduced fat accumulation. In contrast, KO mice failed to respond to pyruvate treatment and did not show any reduction in liver size (**Fig. 6A**).

**Figure 6:**
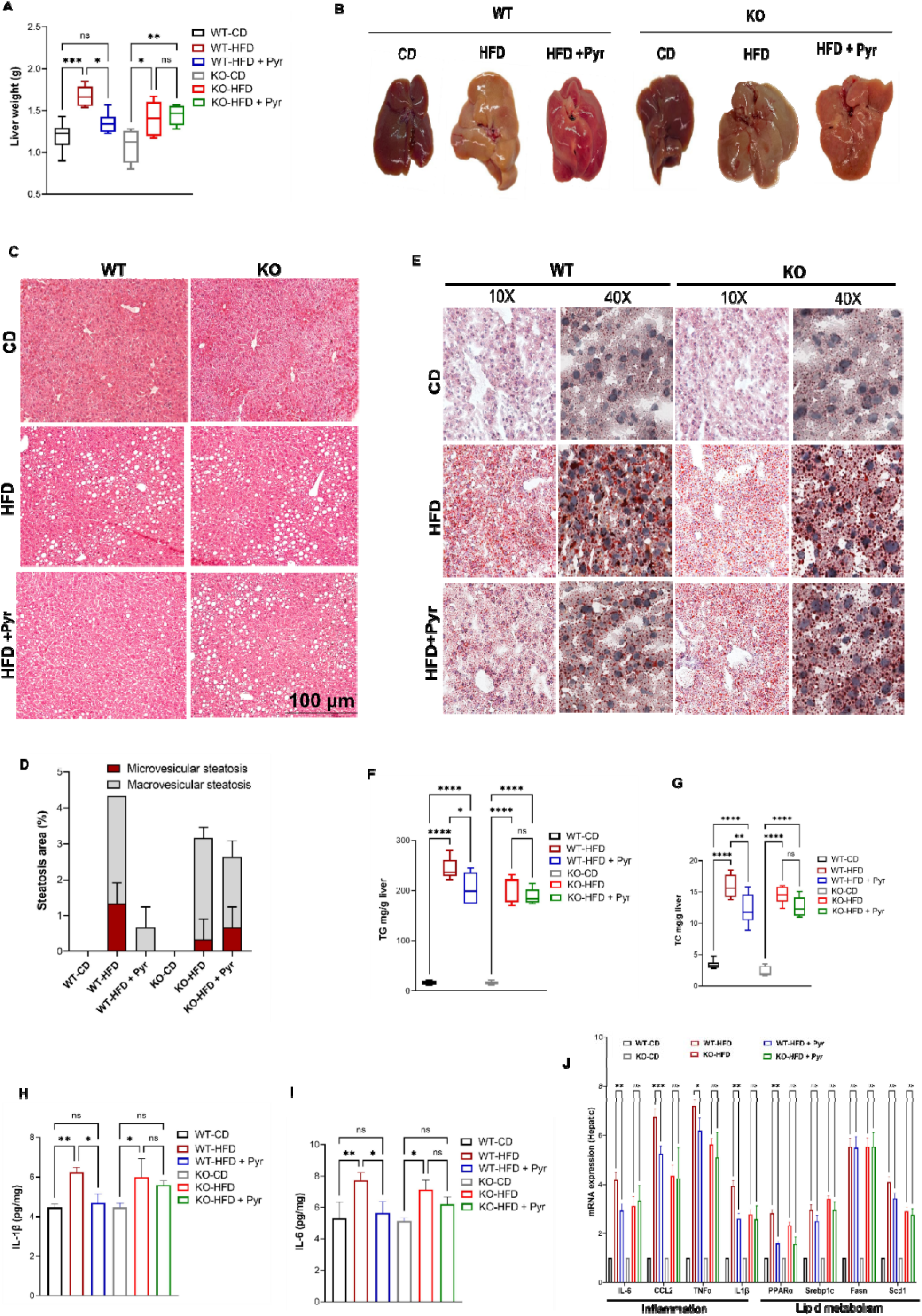
Ablation of cPLA2 impairs pyruvate-mediated dyslipidemia amelioration. cPLA2 WT and cPLA2 KO mice were fed a CD or HFD for 16 weeks with or without pyruvate intervention in drinking water for the assessment of the following. **A.** Weight of liver tissue. **B.** Gross morphologic changes in the liver. **C.** Representative micrographs showing microscopic changes by H&E staining on liver sections. **D.** Quantification of the lipid droplets (steatosis) for each group in panel C. **E.** Representative micrographs showing Oil Red O staining for liver sections showing lipid accumulation. **F.** Hepatic triglyceride level. **G.** Hepatic total cholesterol level. **H-I**. Hepatic mRNA expression of inflammatory genes H. IL-1β and I. IL-6. **J.** Quantitative RT-PCR of the mRNA levels of inflammation and lipid metabolism markers in the liver, as indicated. The expression of WT and KO CD-fed mice, normalized against *Gapdh*, is regarded as 1. Data are mean ± SEM; * p < 0.05, ** p < 0.01, *** p < 0.001. (n=6). Scale bar, 100 μm.

While CD-fed mice showed a healthy reddish-brown liver, HFD-fed mice displayed gross enlargement due to an increase in liver weight. There was a generalized mottled appearance to the livers, which were consistently pale and friable. This pale color signified a prominent deposition of fat within the liver, also known as “marbling.” Pyruvate administration significantly reduced the marbling, as well as the enlargement, indicative of a lean phenotype. In contrast, pyruvate supplementation had no effect on liver weight and marbling in mice with a KO genotype (**Fig. 6B**).

Next, we investigated liver histology by employing (Hematoxylin and eosin) H & E staining to estimate the degree of hepatic steatosis in HFD mice. The livers, following 16 weeks of HFD feeding, displayed markedly increased fat accumulation characterized by vacuolation (exhibiting both macrovesicular and microvesicular steatosis) compared with mice fed normal CD. However, in WT mice, pyruvate administration demonstrated protective effects against HFD-induced hepatic steatosis by alleviating these histological alterations. In contrast, pyruvate-mediated amelioration of hepatic steatosis became ineffective in cPLA2 ablated group, where the vacuolation between the HFD-fed group with or without pyruvate was comparable **(Fig. 6C-D).**

According to the results of Oil Red O staining of liver sections from HFD-fed mice, both WT and KO mice showed an increase in neutral lipid droplets, although it was more prominent in WT mice, whereas CD-fed mice exhibited normal histology. Pyruvate-treated mice displayed noticeably reduced staining, indicating lower fat accumulation. However, in cPLA2 knockout mice, this mitigation was not observed, and fat accumulation was comparable between both groups (**Fig. 6E**).

The hepatic lipid profile validated these results and indicated that HFD induced a significant increase in lipid content characterized by higher levels of TG and TC in both WT and KO mice. The pyruvate group attenuated the HFD-induced TG and TC levels in WT mice but did not mitigate these levels in cPLA2 ablated mice, where both HFD groups showed elevated lipid content (**Fig. 6F-G**). Hence, both H&E and Oil Red O staining results confirmed that mice fed on HFD were showing elevated lipid accumulation and dyslipidemia, which was mitigated by pyruvate in WT but not in KO mice.

Additionally, the mRNA levels of hepatic inflammatory cytokines and lipid metabolism-associated markers were quantified. HFD feeding led to increased expressions of hepatic inflammatory cytokines (IL-6, CCL2, TNFα, and IL-1β) along with expression levels of lipid metabolism-associated markers (PPARα, Srebp1c, Fasn, and Scd1) irrespective of genotype. Pyruvate dramatically inhibited the expressions of inflammatory cytokines improving the liver’s inflammatory profile in WT mice. Pyruvate also repressed the expression of PPARα which is responsible for fatty acid β-oxidation, although, it could not attenuate the expressions of other markers implicated in lipid metabolism even in WT mice. We also determined inflammatory cytokines in the liver lysate by ELISA, which showed higher levels of IL-6and IL-1β in both genotypes. Their levels were significantly attenuated by pyruvate in WT mice but not in cPLA2-ablated mice (**Fig. 6H-I**). In addition these results were largely recapitulated in KO mice where pyruvate could not further mitigate the HFD-induced inflammatory as well as lipid metabolism markers (**Fig. 6J**).

It is imperative to note that obesity is not only associated with fat deposition in the liver but is also often accompanied by ectopic fat deposition in other organs. As a means of evaluating structural changes, we performed H & E staining of the spleen, kidney, and heart isolated from different experimental groups. There was an increase in the ratio of white pulp to the red pulp in the spleen of obese mice, as evidenced by the absence of demarcation between the two types of pulp. There was also an increase in lymphocyte infiltration in HFD experimental groups compared to the control group. As a result of lipid accumulation in the spleen sections of both WT and KO groups following HFD feeding, sinusoidal dilations were also observed. Due to decreased fat accumulation by pyruvate in WT mice, these phenotypes were reverted to being lean mice with normal histology, restoring the white-to-red pulp ratio and decreasing sinusoidal dilation. Conversely, in cPLA2 KO mice, pyruvate failed to reduce histological damage to the spleen and to alter the obliterated structure of the splenic microvasculature (**Fig S7A-B**). Moreover, the weight of the spleen remained non-significantly different between all experimental groups (**Fig. S7C**) with mild hyperemia detected in the HFD-fed groups in both genotypes (data not shown).

Histological analysis of kidneys in all experimental groups revealed well-preserved parenchyma and normal podocytes. In obese mice, the glomeruli exhibited an increased Bowman’s capsule area with an enlarged Bowman’s space as evidenced in the representative histological slide, in both genotypes. In WT mice, the pyruvate-supplemented group did have normal histology, with Bowman’s capsule area comparable to lean mice (**Fig. S7D-E**). The kidneys exhibited a slight increase in weight, but it did not reach significance in obese mice, and all the experimental groups displayed comparable values (**Fig. S7F**).

Obese mice in both genotypes displayed increased cardiomyocyte cross-sectional area and thickened cardiac fibers when compared with respective control mice. The increased cardiomyocyte area was not affected by pyruvate administration in either WT or KO HFD-fed mice (**Fig. S7G-H**). There was, however, a significant increase in the size of the hearts in obese mice, which indicated cardiomegaly (data not shown). This was validated by the observation that the hearts of obese mice were heavier than those of control mice in both genotypes. Even though pyruvate did not influence the cardiomyocyte area, mice administered pyruvate showed a reduction in weight. KO mice, however, did not show a significant difference in weight between HFD and HFD administered with pyruvate (**Fig. S7I**).

### cPLA2 deletion abolishes pyruvate-mediated suppression of inflammation and adipogenesis *in vitro*

To determine the significance of cPLA2 in mediating pyruvate’s anti-inflammatory *in vitro*, we isolated Bone marrow cells from WT and KO mice from different experimental groups and differentiated them into macrophages. These BMDMs were treated with TNFα with/without low (2mM) and high concentrations (4mM) of pyruvate. The qRT-PCR and ELISA results demonstrated that pyruvate dose-dependently ameliorated TNFα-induced mRNA expressions and releases of IL-1β and IL-6, respectively, in cPLA2 WT cells; however, this inhibition was entirely lost in the cPLA2 KO cells (**Fig. 7A-D**).

**Figure 7:**
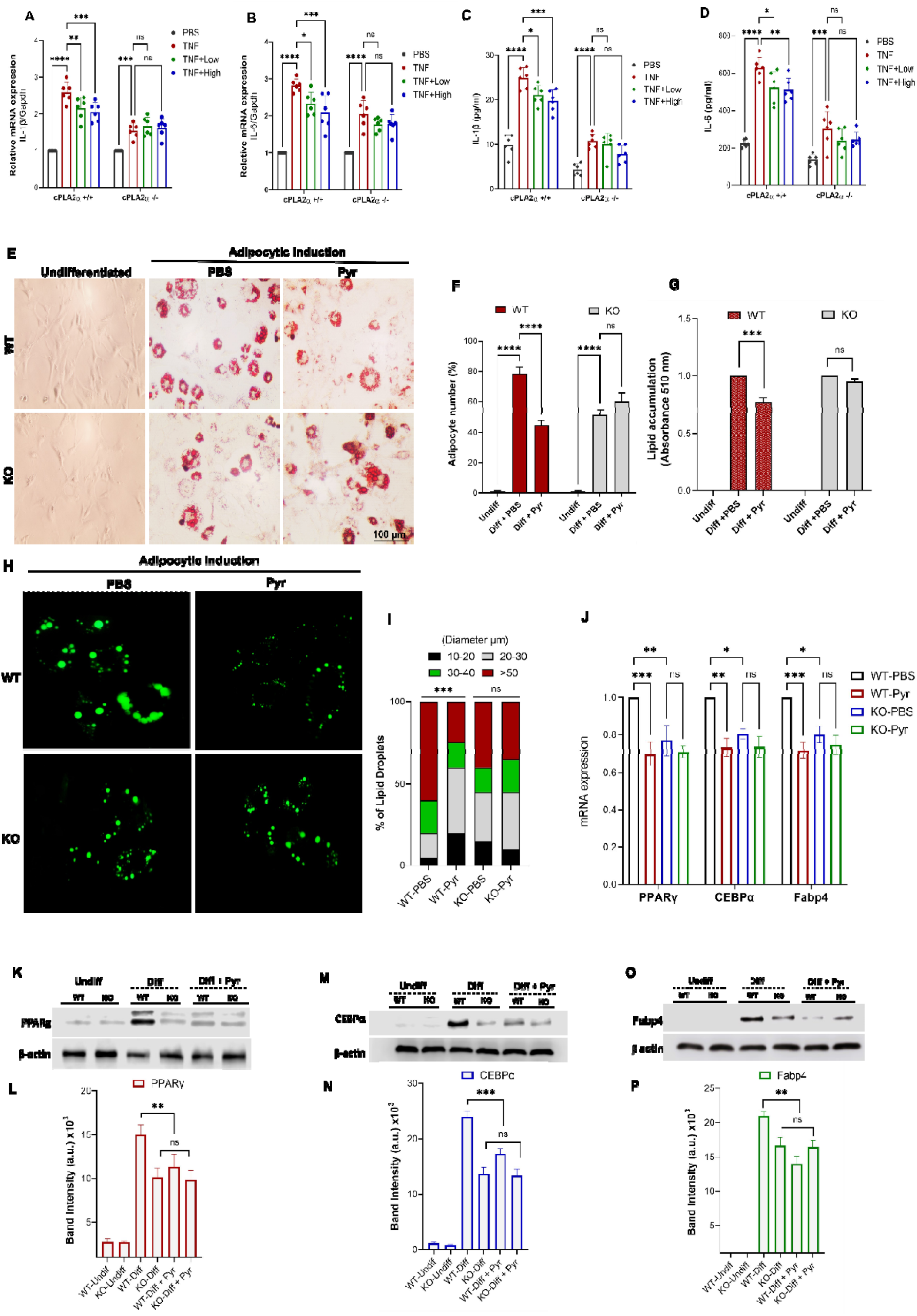
Loss of cPLA2 impairs pyruvate-mediated inflammation and adipogenesis *in vitro*. **A-D.** Bone marrow cells from cPLA2 WT and KO mice from different experimental groups differentiated into macrophages. These BMMLJ were treated with TNFα with/without low (2mM) and high concentrations (4mM) of pyruvate. A-B. The mRNA expression of IL-1β and IL-6 was tested by qPCR. B-D. The secretory level of IL-1β and IL-6 was detected by ELISA. **E-G**. E. Representative images of Oil Red O (ORO) stained lipid droplets (stained red) in preadipocytes induced with adipogenic differentiation with or without pyruvate for eight days, from cPLA2 WT and KO mice. F. Quantification for adipocyte number in WT and KO groups with or without pyruvate. G. Lipid accumulation measured by the colorimetric quantification of the ORO release from indicated groups. **H-I.** H. Microphotographs of lipid droplets of *in vitro* differentiated preadipocytes for indicated groups stained (green) with fluorescent lipid dye. I. Droplets were measured and divided into four size groups: 10-20, 20-30, 30-40, and >50 µm. The Size and distribution of lipid droplets were measured by Image J software. **J.** Expression of PPARγ, C/EBPα, and Fabp4 genes by real-time qRT-PCR in differentiating preadipocyte cells from cPLA2 WT and KO mice, untreated or treated with pyruvate. All values were normalized with respect to β-Actin. **K-P.** Immunoblot analysis of homogenate from *in vitro* differentiated preadipocytes on day eight, for the indicated groups. β-Actin was used as a loading control. K. PPARγ protein level. L. Quantification of panel K. M. C/EBPα protein level. N. Quantification of panel M. O. Fabp4 protein level P. Quantification of panel O.

Since adipogenesis is an irrefutable anti-obesity target, we next evaluated the role of cPLA2 in pyruvate-mediated suppression of adipogenic differentiation. Therefore, we harvested preadipocytes from WT and KO mice and induced adipogenic differentiation with or without pyruvate for eight days followed by assessing their adipogenic phenotypes. Oil O red staining was used to determine the degree of adipogenic differentiation and for quantifying the abundance of lipid droplets (**Fig. 7E**). A diminution in size and the number of lipid vacuoles were noted in cells treated with pyruvate from cPLA2 WT mice. The colorimetric quantification of the ORO stain also confirmed a marked decline in elution from adipocytes supplemented with pyruvate. However, adipogenic differentiation was weaker in KO cells when compared to WT cells, but the treated group displayed no further reduction either in the number of lipid vacuoles or the elution of the oil-red staining (**Fig. 7F-G**). Consequently, cPLA2 is indispensable for pyruvate-mediated suppression of adipogenesis.

Following Oil red O (ORO) staining, fluorescent staining was used as another method to visualize lipid droplets (LDs). This was accomplished by isolating and differentiating primary preadipocytes into adipocytes from each genotype, with or without pyruvate, followed by fluorescent staining. Adipocytic induction in the WT group resulted in an increase in triglyceride accumulation, resulting in the enlargement of lipid droplets. Additionally, the number of LDs in this group was significantly greater than that in the pyruvate-treated group, where both the size and the number of LDs were significantly decreased. There was a general decrease in the LD of adipocytes from KO mice compared with those from WT mice. It should be noted that pyruvate did not further decrease the number in this group, and the incidence of LD diameter remained similar between the treated and untreated KO groups (**Fig. 7H-I**).

mRNAs were extracted from these cells and expressions of the adipogenic factors including PPARg, C/EBPa, and Fabp4 were also analyzed. In both WT and KO cells, these markers demonstrated elevated levels during differentiation. However, there was a substantial difference in the expressions of these markers between WT and KO cells. Despite this, the pyruvate-treated cells displayed considerably reduced levels of PPARg, C/EBPa, and Fabp4 in WT cells, whereas, in KO cells, the mRNA level of these genes did not significantly differ in cells differentiated with or without pyruvate (**Fig. 7J**). Additionally, immunoblot was performed on proteins extracted from differentiating cells. The protein levels of PPARg (**Fig. 7K-L**), C/EBPa (**Fig. 7M-N**), and Fabp4 (**Fig. 7O-P**), were strongly reduced in pyruvate-treated cells compared with PBS-treated WT cells. Nevertheless, the levels of proteins in KO cells were not affected by pyruvate. Taken together, pyruvate inhibits the process of differentiation into mature adipocytes in a cPLA2-dependent manner.

## Discussion

In recent years, pyruvate has gained considerable interest as a dietary supplementation and a potential therapeutic agent, and several studies have demonstrated its role in an array of pathophysiological conditions, including body weight loss, which was first reported over two decades ago [44]. Despite these findings, its functional role and molecular mechanism in HFD-induced obesity have not been distinctly identified and remain largely obscure.

Obesity, a state of chronic low-grade systemic inflammation leading to metabolic inflammation (or meta-inflammation), has been classified as another pandemic due to its substantial rise in global prevalence [45]. Adipogenesis is a complex process in which preadipocytes mature into lipid-bearing adipocytes. It is a crucial process in the determination of the number of adipocytes, making it an imperative therapeutic target for obesity management.

Here we have shown that dietary pyruvate attenuated adipogenesis and downregulated the expressions of the comprehensive network including key transcription factors (TFs) of adipogenesis viz., C/EBPα and PPARγ [46]. These TFs are also responsible for the expression of key proteins that induce mature adipocyte formation like FABP4. Our results indicated that pyruvate mitigated the expression of FABP4 confirming its inhibitory effect on adipocyte maturation [47].

Animals fed a high-fat diet have been shown to develop obesity and metabolic syndrome, as well as its manifestations, including WAT inflammation [48], metabolic dysregulation, hepatic steatosis, and NAFLD, thus mimicking different physiological and metabolic obese phenotype [49]. Consequently, we examined the effect of pyruvate on metabolic syndrome using HFD induced obesity model. By the end of ten weeks, successful induction of obesity was confirmed by increased body weight, adipocyte cell death following adipose tissue expansion, ATM infiltration, liver fat accumulation, hyperglycemia, and other pathological conditions. We observed an increase in body weight and as well as in abdominal circumference, due to hypertrophy and hyperplasia of the WAT at the endpoint of HFD feeding [50]. The results of our study demonstrated that pyruvate was effective in attenuating the dysmetabolic state in part because visceral adiposity is strongly correlated with this phenotypic state [51]. Of note, the results showed that upon HFD feeding, females accumulated comparable vWAT and sWAT fat depots, unlike male mice where vWAT was significantly higher than sWAT in both CD– and HFD-fed conditions. These sex-dependent findings w.r.t WAT were consistent with previous reports [52]. However, despite the precise anatomic location of these depots that differ in mice from humans, these mice depots impact metabolism similar to human vWAT [53]. Furthermore, the mice fed with HFD also showed the presence of chronic inflammation evidenced by elevated basal levels of circulating proinflammatory cytokines. Nevertheless, pyruvate administration reduced the diet-induced weight gain and alleviated the inflammatory status in obesity by reducing circulating inflammatory molecules.

Obesity is a condition referring to increased WAT mass. Besides serving as an energy reservoir, WAT is recognized as a key regulator of metabolic pathways including inflammation, which has been associated with obesity and adipose tissue dysfunction [54]. In our experiments, obese mice displayed WAT inflammation, which was evident by a greater incidence of adipocyte death and the recruitment of macrophages characterized by crown-like structures as apparent from histology. These CLS are a prominent characteristic of low-grade inflammation in WAT, and their density corresponds with the degree of obesity [55]. The results indicated that pyruvate supplementation significantly reduced adipose tissue macrophages (ATMs) and subsequent CLS formation compared to the obese WAT, recuperating in the direction of optimal WAT function.

During obesity, ATMs are known to experience dramatic variations in both humans and rodents [54] and secrete a number of proinflammatory cytokines that result in both local and systemic inflammation [56]. Therefore, ATMs are recognized as viable targets for treating chronic inflammation and obesity-related metabolic disorders. But as of this point, the pharmaceutical agents that target ATMs to treat obesity are inadequate [57]. In this study, we observed predominantly anti-inflammatory M2 macrophage subpopulations in CD-fed lean mice, whereas HFD-induced obese mice showed an elevated infiltration of proinflammatory M1 macrophages which resulted in the release of inflammatory cytokines such as IL-6 and IL-1β [58]. The administration of pyruvate was able to reverse the increased M1 to M2 ratio by driving macrophage polarization toward M2.

NF-kB signaling unites inflammatory and metabolic responses and hence is widely recognized as a crucial factor contributing to obesity and associated metabolic diseases [59]. Considering its pivotal role in all inflammatory processes related to obesity and metabolic disorders, the NF-κB signaling pathway has become another potential target for pharmacological intervention. As it is established that HFD increases NF-kB signaling in mice [60], we employed an *in vivo* non-invasive imaging technique in reporter mice for analyzing the NF-kB signaling in obese mice administered with pyruvate, and revealed that pyruvate administration substantially mitigated the NF-kB signaling in comparison to the untreated obese mice.

Based on our findings, pyruvate suppressed the TNFα induced nuclear translocation of p65. It was reported that NF-κB directly promotes adipogenesis [61], hence, the attenuation of p65 translocation and following NF-kB signaling also explains the inhibition of adipogenic differentiation of preadipocytes as well as the decreases in the expression of adipogenic genes, including PPARγ and CEBPα, in adipose SVF cells by pyruvate. This is also in agreement with a report showing that pharmacological restriction of the NF-κB pathway attenuated the differentiation of 3T3-L1 preadipocytes and human adipose stem cells into adipocytes *in vitro* [62].

The extensive use of HFD-fed mice has undoubtedly helped us to gain a deeper understanding of the mechanisms that regulate glycemic control in obesity [63]. Accumulating evidence suggested that metabolic low-grade inflammation in WAT might play an essential role in the development of obesity-related glucose dysfunction, resulting in an increase in total serum cholesterol and LDL-C concentrations, and is associated with dyslipidemia [64]. Additionally, proinflammatory cytokines are also known to regulate cholesterol metabolism which may in turn alter lipid metabolism. [65]. In our results, clearly as compared to the lower blood glucose levels in lean mice, HFD mice showed elevated glucose levels resulting from dysregulation in the glucose metabolism and displayed dyslipidemia with an elevated lipid profile. Pyruvate, however, was able to regulate this glucose instability in parts and mitigated the overall picture of dyslipidemia.

There is a widespread upregulation of pro-inflammatory signaling cascades in obesity including mediators like IL-6 and TNFα [66]. A study from Kolak et al., [67] confirmed that obese patients with inflamed WAT have more liver fat than subjects without WAT inflammation. As mentioned above, the mediators participate in well-orchestrated and strictly regulated signaling cascades and their disruption may account for systemic low-grade inflammation-mediated interaction between obesity and NAFLD. NAFLD is manifested by excessive hepatic fat deposition caused by carbohydrate and/or lipid intake. In our study, the HFD-fed group presented higher fat accumulation (steatosis) in the liver tissue with elevated hepatic lipids as compared to the lean mice, which is consistent with other studies [68]. Pyruvate showed significantly alleviated fat deposition in the liver as well as reduced chronic lipid overload as compared to HFD mice, thereby controlling dyslipidemia. But the mRNA expression of different hepatic genes asserted that the inhibition in the overall NAFLD by pyruvate was not due to the decreased expression of lipogenic genes including SREBP-1c, FASN, and SCD-1 but due to the attenuation of the inflammatory markers including the secretory level of IL-1β and IL-6.

Combined use of several approaches, including DARTS, proteomics, CETSA, and ITDR, led to the isolation of cPLA2α as a previously unidentified target of pyruvate. After identifying cPLA2 (α type) as the target of pyruvate, we established the HFD obesity model in cPLA2 WT and KO mice. As shown in C57BL/6 mice, pyruvate replicated its protective effects by effectively preventing obesity and metabolic syndrome in response to a high-fat diet in a cPLA2 WT mice manner. We further conducted a loss-of-function analysis to confirm that cPLA2 is molecular target of pyruvate, by ablating cPLA2 globally in mice and inducing HFD obesity. The results revealed that the deletion of cPLA2 abrogated the protective effects of pyruvate on obesity. Hence, confirming that pyruvate targets cPLA2 to mediate the regulation of metabolic syndrome including adiposity, meta-inflammation, and NAFLD in HFD-induced obesity.

Intriguingly, cPLA2 mediated production of AA is also known to activates NF-ĸB signaling in different tissues and plays a role in the pathogenesis of inflammatory diseases [69]. However, it should be noted that the contribution of cPLA2, being a canonical regulator of AA metabolism, to eicosanoid synthesis producing variety of lipid mediators, is relatively insignificant in WAT [70]. This may explain why pyruvate selectively mitigated the mRNA expressions of inflammatory genes in WAT, but not lipogenic genes. Based on these findings, it can be concluded that pyruvate functions in a cPLA2-dependent manner and that its mechanism of action is primarily dependent on the cPLA2-AA-NF-ĸB axis in attenuating meta-inflammation, obesity and other obesity-mediated complications [71].

In conclusion, this study is the first to provide molecular insights into the mechanisms through which pyruvate alleviates high-fat diet**-**induced obesity and accompanying metabolic syndrome. It identified cPLA2 as a novel target of pyruvate, providing a comprehensive mechanistic insight into the role of pyruvate in obesity. Moreover, it provides the groundwork for further investigations into pyruvate/cPLA2 interaction. As pyruvate functions as a potent antagonist of cPLA2, it can be used to potentially treat a variety of cPLA2 driven diseases. Accordingly, these findings could have implications for the development of novel dietary supplementations and/or therapeutics to benefit a complex disease like obesity, extending the therapeutic role of pyruvate beyond its use as an over-the-counter energy supplement (OTC).

## Material and methods

### Identification and characterization of the binding of pyruvate to cPLA2

Drug affinity responsive target stability assay (DARTS), proteomics, CETSA, and mass spectrometry were used.

## Ethics approval

All animal procedures were carried out in accordance with institutional guidelines and approved by the Institutional Animal Care and Use Committee of New York University.

## Isolation and differentiation of preadipocytes

Preadipocyte isolation was performed according to the protocol of a commercial preadipocyte isolation kit (ab 196988) kit.

## Acknowledgments

We are thankful to Proteomics Laboratory for protein-related mass spectrometry analysis and the preclinical Imaging Laboratory at NYU Grossman School of Medicine.

## Conflict of interests

The authors declare no competing interests.

## Funding

This work was partially supported by NIH research grants R01AR062207, R01AR061484, R01AR076900, R01AR078035 and R01NS070328.

## Author Contributions

S.H. and C.J.L. conceived, designed, and directed the study. SH performed most of the experiment with the assistance of N.G (blinded observer for all histology scorings). X.Z (Blood collection from C57BL/6 mice); J.G. (H & E of colons). A.H, H.L.W, J.L (dissections, embedding for histology) S.H., N.G., and C.J.L. wrote the manuscript with inputs from all authors.

## Notes

### Competing Interest Statement

The authors have declared no competing interest.

