## Supplementary material for "Dietary Pyruvate Targets Cytosolic Phospholipase A2 to Mitigate Inflammation and Obesity in Mice"

This file includes:

**Material and methods (Pages 2-14)**

**Figures S1-S7 (Pages 15-27)**

**Table S1 (Page 28)**

### Material and methods

#### Mice

All animals were housed on a 12-hour light-dark cycle with ad libitum access to food and water in a specific pathogen-free environment. Animal protocols were approved by the Institutional Animal Care and Use Committee (IACUC) of New York University Grossman School of Medicine. In all cases, unless stated otherwise, mice were male, typically 8 weeks old, and age-matched. C57BL/6J background, NF-κB Luc, TNFα-transgenic mice (TNFα-tg), and cPLA2 were the genotypes used for the experimental studies. C57BL/6J and NF-κB- luciferase reporter (model number 10499) mice were purchased from Jackson Laboratory (Bar Harbor, ME, USA). cPLA2 (+/-) heterozygous mice were a generous contribution from Dr. Naikui Liu's lab at Indiana University School of Medicine. These mice originated from Joseph V Bonventre's lab at Brigham and Women's Hospital. These cPLA2 mice are on a C57BL/6 and 129 mixed backgrounds. A heterozygous cPLA2 (cPLA2 +/-) mouse was crossed to generate wild-type littermates (cPLA2 WT) and a global knockout (cPLA2 KO) mouse. TNFα-tg mice were generated by transfecting the human TNFα gene to C57BL/6 background mice, which mimics a systemic inflammatory state [1].

#### High-fat diet (HFD) induced obesity mouse model.

All mouse high-fat or chow diet feeding experiments were performed with age-matched littermate controls which were males unless stated otherwise. Prior to the start of the experiment, mice were adapted to the modified conditions for 1 week. In the meantime, mice were distributed into four groups (n = 8-10 per cage) in C57BL/6 (both male and female) and NF-κB Luc mice: the chow diet fed group (CD), high-fat diet group (HFD), the high fat-diet group supplemented with 1% pyruvate in drinking water (HFD + Pyr) and high fat-diet group administered with 10 or 20 mg/kg orlistat orally (HFD+ Orli) as a positive control. At eight weeks of age, mice were switched to high fat-diet (Diet, D12492i) with 60% kcal from fat, 20% kcal from carbohydrate, and 20% kcal from protein for 10 weeks. For cPLA2 WT and KO mice, the division of groups for each genotype was the same (CD, HFD, and HFD

+ Pyr) as in C57BL/6 mice however, the experimental feeding time was 16 weeks. Pyruvate-containing drinking water was freshly prepared and replaced every two days. Bodyweight and food intake were recorded every week and they were maintained.

**Blood and tissue isolation**

At the end of the experimental period, all animals fasted for 12 h, anesthetized using 200 μl anesthetic agent containing 20 mg/ml ketamine and 2 mg/ml xylazine followed by whole blood collection from each mouse were placed on ice unless stated otherwise, and finally euthanized using $CO_2$ confirmed by cervical dislocation. Multiple tissues including the liver, heart, spleen, kidneys, subcutaneous white adipose tissues (sWAT), and visceral white adipose tissue (vWAT) were harvested, rinsed, weighed, either embedded or snap frozen in dry ice, and stored at -80 °C until further processing. Serum was collected from whole blood after centrifugation at 2000 × g for 20 min at 4 °C and stored at -80 °C until further analysis. Adipose tissue and liver tissues were homogenized with lysed following the kits' instructions (NBP2-37853). These tissue lysates and plasma were used for biochemical and histological analyses [2].

**Oral Glucose Tolerance Test (OGTT)**

A glucose tolerance test was performed in mice after an overnight fast. The glucose concentrations were determined with a blood glucose meter (Accu-Check Active 1, Roche Pharmaceutical Ltd., Basel, Switzerland) and measured in blood collected from the

tail vein immediately before and, 15, 30, 45, 60, and 120 min, after oral administration of glucose at 2 g/kg [3]

**Mouse Genomic DNA isolation**

Mice tails of size ~4 mm was snipped and collected in sterile 1.5 ml microcentrifuge tubes. It was followed by the addition of 100 μl of a tissue digestion buffer (NaOH 25 mM and EDTA 0.2 mM) to each sample. These samples were then heated at 100 °C in a heating block (Isotemp, Fisher Scientific) for 1 hour. After thermal digestion, neutralization buffer (100 µl of 40 mM Tris buffer, pH 7.5) was added to each tube. The tubes were then centrifuged for pelleting tail debris at 14,000 g for 10 minutes. The genomic DNA concentration in the supernatant was measured using a spectrophotometer (Nanodrop 200 C, Thermo Scientific NanoDrop 2000). DNA concentration was tweaked to 50 ng/µl by adding nuclease-free water. Finally, the isolated DNA samples were stored at −20 °C in microcentrifuge tubes until further use.

**Genotyping**

Genotyping was performed on genomic DNA from tail biopsies by PCR before proceeding with experiments [4]. For PCR amplification, GoTaq green master-mix (10 µl, Promega, M7123) and 1 µL of each of the forward and reverse primers from 100 µM stock (Integrated DNA Technologies, USA) pre-diluted to 10 µM were added in each 0.2 ml PCR tube. Finally, 5 µl of the previously isolated genomic DNA sample and 2 or 3 µl of nuclease-free water were added to each tube to obtain a final reaction volume of 20 µl. PCR was performed using a standard thermocycler (SimpliAmp™ Thermal Cycler). The amplification conditions consisted of an initial denaturation (95 °C for 3 minutes), followed by 34 cycles of denaturation (94 °C for 30 secs), annealing (56-59 °C for 30 secs), and extension (72 °C for 30 secs) and a final extension at 72 °C for 4 minutes. PCR products were removed from the thermocycler and maintained at room temperature for a few minutes and then were analyzing analyzed by 1.5% (w/v) agarose gel electrophoresis with ethidium bromide [1].

For cPLA2, PCR genotyping required three primers, one reverse primer (cPLA 604, 5’-GACTCATACAGTGCCTTCATCAC-3’) and two forward primers (cPLA 3F, 5’-TGTGTACAATCTTTGTGTTGTTTCA- 3’ and PgkNeo 5’-GGGAACTTCCTGACTAGGGG-3’). The PgkNeo and cPLA 604 primers amplify a 300bp fragment from the knockout allele whereas cPLA 3F and cPLA 604 amplify a 100 bp fragment from the wild-type cPLA2 gene [5]. For the TNFα-tg genotype, one reverse primer (5'- CGGGCCGATTGATCTCAGC-3') and one forward primer (5'-GAGGCCAAGCCCTGGTATG-'3) were used. The presence of TNF transgene was validated by the presence of 91 base pair products. Lastly, for NF-κB Luc mice, one reverse (5'-AGGGTTGGTACTAGCAACGC-3') and one forward (5'-TGGCAGAAGCTATGAAACGA-3') primer were used. The presence of the luciferase reporter gene was evidenced via a 182 base pair product [6].

**Preparation of agarose gels and electrophoresis of PCR products**

Agarose gels (1.5 %) were prepared using agarose of 95% purity (Sigma-Aldrich, A5093) dissolved in 1X Tris-Acetate EDTA buffer (40 mM Tris-acetate, 1mM EDTA, pH 8.3, Fisher Bioreagents BP13324) by heating the solution in an on a hot plate or microwave oven for 3–4 minutes. The agarose solution was kept at room temperature for about 5 min and then 5 µl of ethidium bromide (10 mg/ml, E7637, Sigma Aldrich) was added to the melted agarose and was immediately poured on a transparent gel casting tray (Bio-Rad PowerPac Basic w/ Wide Mini Sub Cell GT Electrophoresis Unit) fitted with a 10 well comb. The gel tray was placed on the electrophoresis system and the electrophoresis chamber was filled with 1X TAE buffer till about 1.5 cm above the gel. 5-8 µl of each PCR product was loaded into each well and electrophoresis was performed for 80 minutes at 6.5 volts/cm. The power pack supply (1000/500 power supply, Bio-Rad) was set to 90 V. A DNA size marker of 100 bp (Gene ruler 1 kB plus, DirectLoad™ Plus 1kb DNA Ladder) was used. After the run was complete, the gel images were acquired using a gel documentation system (Biorad Gel Doc XR Imaging System).

**Histological analysis**

The liver and epididymal adipose tissue from a representative mouse in each group were fixed in 10% buffered formalin, embedded in paraffin, and cut into 8 µm thick sections. Some sections were stained with hematoxylin and eosin (H&E) for histological examination of lipid droplets and images were acquired using an Olympus SZX10 microscope [7].

### Cell culture experiments and treatments

All cells viz., bone marrow cells, bone marrow-derived macrophages (BMMϕ), RAW264.7 cells (obtained from ATCC) were maintained in DMEM (without pyruvate, DMEM, Gibco™ 11995073) supplemented with 10% Fetal Bovine Serum (FBS, Gibco™- 000044) and 1% Penicillin-Streptomycin (P/S Gibco™ 15140122) at 37 °C under 5% CO2 in a humidified incubator. Cells were detached using 0.05% trypsin (Gibco™ 25300054) if sub-culturing was required. The differentiation media were supplemented with different pyruvate (sigma) concentrations viz., 0.02, 0.06, 0.1, 0.2, 2, 4, 8, and 10 mM for screening. After the screening, low (2mM) and/or high (4mM) concentrations of pyruvate were used unless stated otherwise. Mentioning only pyruvate indicate the use of high concentration (4mM). In all the experiments cells were incubated with or without low and high concentrations of pyruvate for 24 hours if the sample is to be collected for real-time PCR or 48 hours if performing ELISA. In the experiments that were subjected to TNFα stimulation, the concentration used for TNFα was 10 ng/ml.

### Nuclear translocation and Immunofluorescence staining

BMMϕs were cultured on coverslips in 12- wells culture plates in the absence or presence of pyruvate (4mM) and were stimulated by TNFα (10ng/mL) followed by 6 hours of incubation. Immunofluorescence was performed to visualize the subcellular localization of p65. After incubation, the cells were fixed with 4% PFA for 10 min at room temperature, then permeabilized in 0.1% Triton X-100 for 10 min, followed by blocked in 1% BSA for 1 h at room temperature. Cells were then incubated with p65 primary antibody (764s, cell Signaling) with dilution 1:50 - 1:100 at 4°C overnight. The next day, cells were washed three times in PBS for 10 min by gentle aspiration, then incubated with anti-Rabbit IgG conjugated with Alexa Fluor® 488 (1:200, A-11008, Invitrogen) at room temperature for 1h. 4',6-diamidino-2-phenylindole (DAPI, 1:1000, D1306, ThermoFisher Scientific) was used to stain the nucleus for 10 min at room temperature. The coverslips were removed and mounted on microscope slides with Kaiser's glycerol gelatin and further analyzed and imaged using Leica TCS SP5 confocal system while Fluorescence intensity was quantified using ImageJ software [8].

***In vivo* Bioluminescence Imaging for NF-kappaB signaling.**

NF-κB Luciferase reporter mice were employed to evaluate NF-κB upon HFD feeding via *in vivo* bioluminescence. For screening, [9]. Mice from different groups of CD, HFD and HFD+ Pyr/Orlistat were imaged (n = 6 per group the end of experimental period. *In Vivo* Imaging System (IVIS) was performed. Mice were anesthetized with isoflurane and an intraperitoneal injection of 150 mg/kg luciferin was administered. Imaging was performed (after 10 minutes) every 5-10 minutes extended over a period of up to 60 minutes to determine the peak signal. Bioluminescent signals were quantified using a whole animal bioluminescence imaging system (PerkinElmer IVIS® Spectrum) and analysis software (PerkinElmer Living Image®) [6].

***In vitro* differentiation of primary bone marrow-derived macrophages (BMMϕ)**

Bone marrow cells were isolated from C57BL/6 and TNFα-tg mice. After mice were sacrificed by cervical dislocation, the tibia and femur were isolated under aseptic conditions and washed in PBS containing antibiotic/antimycotic solution (1%, Invitrogen, USA). Both ends of the bones were cut to open the bone medullary cavity and were placed in a 1.5 mL sterile microtube, bone marrow cells were collected by centrifuge at 13000g for 90 seconds and isolated cells were suspended in Dulbecco's modified Eagle's medium (DMEM) (Gibco, USA) supplemented with 10% fetal bovine serum (FBS; Gibco, USA), and antibiotic/antimycotic (Gibco-Brl #15240-062,) with M-CSF (10ng/ml, 576406, Biolegend) and incubated at 37 °C, 5% $CO_2$ for 6 days to allow differentiation.

**RNA extraction and quantitative Real-time PCR (qRT-PCR)**

Total RNA was extracted from the desired cell or tissue type by using RNeasy Mini Kit (74106, Qiagen or adipose tissue RNA Purification Kit (RNeasy Lipid tissue kit 74808). The quality and concentration of total RNA was determined spectrophotometrically using NanoDrop (Thermoscientific). First-strand complementary DNA (cDNA) synthesis was performed with 1 µg RNA using a High-Capacity cDNA Reverse Transcription Kit (4368813, Applied Biosystems). SYBR green-based (4367659, Applied Biosystems). Gene expression levels were quantified using quantitative PCR in triplicate on the Real-Time PCR System (Applied Biosystems) using target genes specific primers listed in supplementary table 1. Relative mRNA expression was calculated by normalizing the expression level of target genes with the housekeeping genes viz., glyceraldehyde-3-phopshate (*Gapdh*) or *β-actin*. The relative transcription levels of the mRNA were calculated according to the $2^{-\Delta\Delta Ct}$ formula and reported as a relative mRNA fold change [10].

**Immunofluorescence imaging Lipid droplets**

Lipid droplets in differentiated adipocytes were identified with lipid dye droplet fluorescent using LipiDye II (#FNK-FDV-0027) following the manufacturer's protocol. Briefly, differentiation culture medium was removed, and cells were washed with PBS and fixed with 4% paraformaldehyde. The cells were washed three times in PBS and incubated with working concentration (1µM) of LipiDye II-containing medium at 37 °C over 10 minutes. Cells were then washed once with PBS and were imaged using a Nikon Eclipse Ti-U fluorescence microscope (Tokyo, Japan) on a 10x objective lens.

**Oil red O staining**

Primary preadipocytes cells differentiated in adipogenic media for 8 days in the absence or presence of pyruvate, were washed with PBS followed by to Oil Red O staining assay according to the protocol of a commercial kit (Abcam, #133102). Briefly, cells were first washed with PBS and fixed with formalin solution for 15 min. The fixed lipid droplets were then stained with Oil Red O solution for 30 min at room temperature. Microscope images were taken to visualize.

red oil droplets staining in differentiated cells.

**Triglyceride accumulation assay**

Primary preadipocytes cells differentiated in adipogenic media for 8 days in the absence or presence of pyruvate or tissue lysate, were used to determine TG concentrations with a commercial kit (Abcam, #102513). Briefly, total cellular lipids were extracted with lipid extraction solution under heating. The TG content was then determined by adding lipase, which converted TG to glycerol. Glycerol was subsequently reacted to convert the probe to generate color, which can be measured spectrophotometrically at 570 nm in a plate reader. TG concentrations were calculated based upon a standard curve made from TG standards and normalized to total cellular protein content.

**Enzyme-Linked Immunosorbent Assay (ELISA)**

Cell culture or tissue lysate supernatants were collected after 48 hours of treatment (treated with or without pyruvate or 5-ASA). Abcam Protein Extraction Kit (ab270054) was used following the protocol booklet instructions. Cytokine levels of IL-1β (IL-1β 88-7013, Invitrogen) and IL-6 (IL-6: 88706476, Invitrogen) in cell culture supernatants, sera or colonic tissues from murine models were detected by sandwich ELISA according to product specifications in ELISA kit. For serum samples, whole blood collected from each mouse was subjected to centrifugation at 3000 rpm for 10 min.

**Drug affinity responsive target stability (DARTS) assay and mass spectrometry**

DARTS was performed according to previously reported methods [11]. Briefly, RAW264.7 cells were lysed using M-PER™ Mammalian Protein Extraction Reagent (78501, Thermofisher), followed by centrifugation at 18,000X for 10 min at 4°C. The supernatant (cell lysate) was then incubated with 4mM pyruvate or PBS (Thermofisher, Cat. 70011044) for 1 hour at room temperature and was held on a rotator. This mixture was subjected to digestion by pronase (Sigma, Cat. P5147) at room temperature for 15 min. Samples were boiled after adding 5x SDS loading dye for SDS-PAGE or western blot detection. A band with a molecular weight of ~80kDa was found to be protected by pyruvate treatment. This band was excised and then analyzed by mass spectrometry, performed at NYU Proteomics Laboratory. All MS/MS spectra were collected using the following instrument parameters: the resolution of 15,000, AGC target of 1e5, maximum ion time of 120 ms, one microscan, 2 m/z isolation window, fixed first mass of 150 m/z, and NCE of 27. MS/MS spectra were searched against a Uniprot Human database using Sequest within Proteome Discoverer.

**CETSA assay**

RAW264.7 cells were treated with pyruvate for 1 hour at 37°C with 5% $CO_2$. Cells were harvested and were equally divided into multiple sterile eppendorf tubes. Aliquoted cell suspensions were subjected to thermal digestion by using a range of temperatures (40, 43, 46, 49, 52, 55, 58, 61, and 64 °C) for 3 min. Samples were subsequently lysed through 3 repetitive freeze-thaw cycles and finally centrifuged at 20,000 g. The supernatants were collected for western blot analysis. After determining the overall melting behavior of the target protein, isothermal dose response (ITDR) was performed. For ITDR, a series of concentrations (ranging from 0.2 to 2000 μM) of pyruvate was used for the treatment of cells and a constant temperature (55°C) was selected based on CETSA for thermal denaturation. The remaining steps were followed as previously reported [12].

**cPLA2 activity assay.** BMMϕ were cultured and treated with TNF- α(10ng/ml) for 48 hours. The cell lysate was collected and incubated with or without Pyruvate (2 mM and 4mM), for 1 hour. cPLA2 activity of each mixture was detected by ELISA kit (765021, Cayman). Alternatively, BMMϕ cells were cultured and treated with TNF-α (10ng/ml) with or without Pyruvate/ATK for 48 hours. Cell lysate was collected and the cPLA2 activity was detected by PLA2 activity ELISA kit (765021, Cayman).

**Arachidonic acid detection.** BMDMs were treated with TNF-α (10ng/ml) with or without Pyruvate/ATK, the cultured medium was collected and analyzed using an arachidonic acid ELISA kit (NBP2-66372, Novus Biologicals). The result was detected at 450nm by automatic plate reader. ATK was implemented as a positive control.

**Cell viability assay**

Cell viability assay was performed according to the protocol described in MTT Cell Proliferation Assay Kit (Ab211091). Briefly, cells were grown at varying densities (1-5 x106 cells per mL) into a 96 well plate with clear flat bottom and untreated or treated with various concentration of pyruvate (0,0.02,0.2,1,2,3,8,and 10 mM) for 72 hours. The media was aspirated at each time point and 50 µL of serum-free media and 50 µL of MTT Reagent into each well. For background control, 50 µL MTT Reagent + 50 µL cell culture media without cells. The plate was then incubated at 37°C for 3 hours, following by the addition of 150 µL of MTT Solvent into each well. Finally, the plate was wrapped in a foil to prevent light and put on an orbital shaker for 15 minutes. Finally absorbance was read at OD = 590nm.

**Isolation and differentiation of preadipocytes**

Preadipocyte isolation was performed according to the protocol of a commercial preadipocyte isolation kit (ab 196988) kit. Briefly, adipose tissues were excised from the mice and minced with dissecting scissors in a sterile vessel for at least 5 minutes. 1 mL of Collagenase per 0.5 g of tissue was added and the tube was incubated in a heated orbital shaker at 37°C for 30 minutes at 160 rpm followed by adding 9 mL of Collagenase Stop Buffer per 1 mL of Collagenase. The suspension was filtered through Cell Strainer (100 µm) and the centrifuges at 500*g* for 10 minutes. tHe pellet was resuspended in 1 mL of Red Blood Cell Lysis Buffer for 1 minute. For wash, 9 mL of PBS and the cells were filtered through

70 μm Cell Strainer followed by another cycle of centrifugation (500 x*g* for 10 minutes). Remove the supernatant and resuspend cell pellet in 2 mL of preadipocyte medium. Add cells into 1 well of a 6-well plate with complete DMEM media and incubate at 37°C with 5% CO2. They were maintained for 48 h post-confluence before differentiation. Then fresh preadipocyte differentiation medium (C-27437 sigma) was added and changed every 3 days.

**Hematoxylin and eosin staining**

At the time of sacrifice, samples from organs and adipose tissues were fixed in 10% neutrally buffered formalin and was dehydrated using escalating proportions of ethanol/ water (v/v). Tissue samples were embedded inside paraffin for 16 h. Paraffin-embedded samples were sliced at 6 μm in thickness and were placed in xylene. Standard Hematoxylin and Eosin staining of tissue slides was conducted according to the standard protocol [13].

**Body composition analysis.**

Dual energy X-ray absorption (DEXA) scan was used for the whole-body composition analysis using DEXA scanner (Insight, Osteosys/Sciatica, London, ON, Canada). The instrument was calibrated before each scanning session using a Phantom with known bone mineral density according to the manufacture's instruction.

**Immunofluorescence staining of vWAT.**

It was performed as previously described [14]. Briefly, the vWAT frozen sections frozen sections were fixed with paraformaldehyde for 15 minutes at room temperature, washed with PBS and treated with 0.4% Triton X-100 (Sigma) for 15 minutes, at room temperature for permeabilization.

After washing, the tissue was incubated in 3% BSA for blocking at room temperature for one hour. After washing the tissues were incubated with primary antibody (rabbit polyclonal to F4/80 [1:500,

Cat#: ab60343]; mixed with CD206 (Monoclonal Antibody Cat #MA5-16870) or iNOS (Cat #14-5920-82) overnight at 4 °C. They were then washed thrice with PBS. For immunofluorescence double staining, fluorochrome-conjugated secondary antibodies against a donkey anti-rabbit IgG H&L (Alexa Fluor® 488) (1:1000 dilution) (Abcam) and a goat anti-rat IgG H&L (Alexa Fluor® 647) (1:1000 dilution) (Abcam) were added to the sections and co-incubated at room temperature for 1 hour followed by the microscopic imaging.

**Western blot**

After protein cell/tissue lysates were prepared, protein concentrations were determined using BCA protein kit (ThermoFischer Scientific) using bovine serum albumin (BCA) as standard. Samples were separated by SDS-PAGE and were electroblotted into PVDF membrane (millipore) electrophoretically using a wet transfer system. The membrane was blocked using 5% (w/v) non-fat milk in TBST for half an hour at room temperature, followed by probing the membrane with with an antibody (1:1000 dilution) at 4°C overnight and adding a secondary antibody (1:10,000 dilution) at shaking room temperature for 1 hour. The immunogenic bands on the membrane were developed by chemiluminescent (ECL) substrate (Amersham Biosciences, Pittsburgh, PA) and visualized by the GelDoc system. All the antibodies used were from Cell Signaling Technology unless stated otherwise. Antibodies used: *Cell signaling*- NF-KB p65 (#4764), GAPDH (#5174), cPLA2 (#5249), PPARγ (#2435( and FABP4 (#50699), *Invitrogen*: Lamin B1 (#PA5-19468), PLXNB2 (#PA5- 47880), Flag tag antibody (# MA1-91878-HRP), C/EBP alpha (# MA1-825).

**Statistical analyses.**

The experiments were randomized, most investigators were blinded to experimental allocation and outcome analysis. Statistical analysis was performed by GraphPad Prism (version 9.4’ GraphPad Software Inc.). All values are presented as a mean ± standard error of the mean (S.E.M). Statistical significance between two groups was determined using a two-tailed, unpaired Student's t-test. For comparison of more than two groups, ANOVA analysis was used with Bonferroni post-hoc test unless stated otherwise. For qPCR data, statistical analysis was run on $2^{-(\Delta\Delta Ct)}$ or normalized relative quantity (NRQ) values. All data points represent biological replicates, and the representative results were obtained from at least three independent experiments. For *in vivo* experiments, the numbers of mice employed per genotype were 8-10 unless indicated in figure legends.

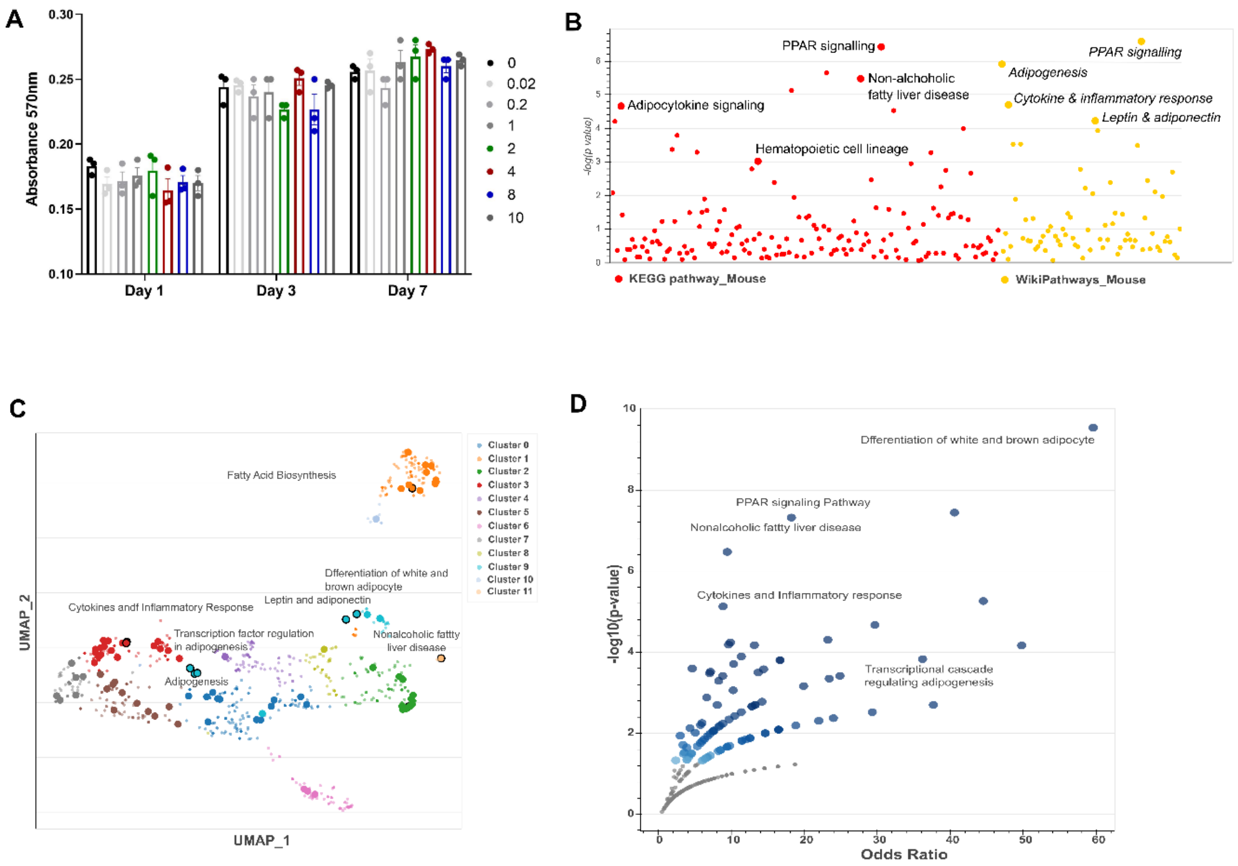


**Figure S1: Bulk RNA seq and pathway analysis revealed inhibition of adipogenic pathway by pyruvate.**

**A.** MTT analysis for the assessment of cell viability.

**B.** Comparison of the KEGG pathway and Wikipathway analysis on the basis of statistical significance (P-value) for differentially expressed genes (DEGs) of bmMSCs treated with pyruvate in the presence (4 mM) in the presence of TNFα for 24 hours. Red dots represent KEGG pathway DEGs and yellow dots represent Wikipathway DEGs.

**C.** Scatterplot of all terms in the WikiPathway gene set library**.** Each point represents a term in the library. An enriched term is more significantly characterized by darker and larger points.

**D.** Volcano plot of terms from the WikiPathway gene set**.** Each point signifies a single term, plotted by the corresponding odds ratio (x-position) and -log10 (p-value) (y-position) from the enrichment results of the input query gene set. A larger and darker point indicates that the input gene set is significantly enriched for the term.

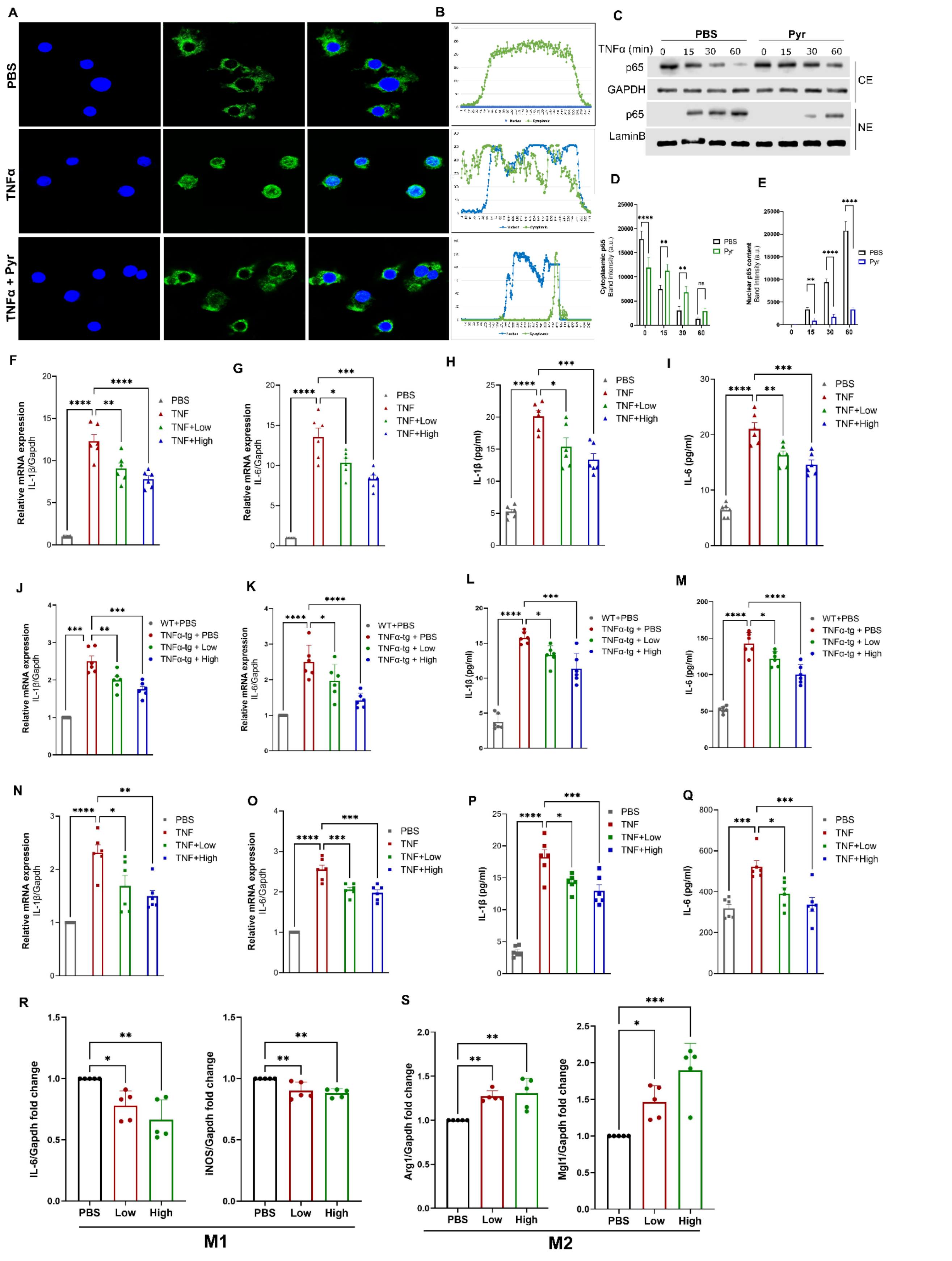

**Figure S2: Pyruvate inhibits TNFα induced NF-κB nuclear translocation and regulates macrophage polarization.**

**A-B.** A. RAW 264.7 cells were cultured with TNF-α in the absence or presence of pyruvate for 6 hours. Immunofluorescence cell staining was performed to visualize the subcellular localization of p65. 4′,6-diamidino-2-phenylindole (DAPI) was used to stain the nucleus. B. Quantification of the translocation, where blue dots represent the nuclear fraction and green dots represent the cytosolic fraction.

**C-E.** C. RAW 264.7 cells were treated with TNF-α in the absence or presence of pyruvate for indicated time points. Cytoplasmic extraction (CE) and nuclear extraction (NE) were examined by performing a western blot with an anti-p65 antibody. Gapdh protein and Lamin B protein expressions served as an internal control for cytoplasmic fraction and nuclear extracts respectively. Quantification using image J for D. cytoplasmic p65 and E. nuclear fraction.

**F-I**. RAW 264.7 cells were treated with TNF-α in the absence or presence of low (2 mM) and high (4 mM) doses of pyruvate for 24 hours for real-time PCR or for 48 hours for ELISA. F-G. The mRNA level of IL-1β and IL-6 were detected by qRT-PCR. H-I. The secretion level of IL-1β and IL-6 was tested by ELISA.

**J-M.** *In vitro* differentiated BMMϕ from TNF-tg mice were treated with a low and high concentration of pyruvate for 24 hours for real-time PCR or for 48 hours for ELISA. J-K. The mRNA level of IL-1β and IL-6 were detected by qRT-PCR. L-M. The secretion level of IL-1β and IL-6 was tested by ELISA.

**N-Q.** *In vitro* differentiated BMMϕ from C57BL/6 mice were treated with TNF-α in the absence or presence of low and high doses of pyruvate with a low and high concentration of pyruvate for 24 hours for real-time PCR or for 48 hours for ELISA. N-O. The mRNA level of IL-1β and IL-6 were detected by qRT-PCR. P-Q. The secretion level of IL-1β and IL-6 was tested by ELISA.

**R-S.** Bone marrow cells were treated with M-CSF, IFN-γ (25ng/ml), and LPS (250ng/ml) or IL-4 (20ng/ml) for type 1 macrophages (M1) or type 2 macrophages (M2) respectively. Low and high dose of pyruvate was used to treat cells, as indicated followed by qPCR analysis of R. *IL-6 and iNos2* mRNA expression in BMMϕ polarized to M1 macrophages and S. *Arg1 or Mgl1* mRNA expression in BMMϕ polarized to M2 macrophages. Data are mean ± SEM; * $p < 0.05$, ** $p < 0.01$, *** $p < 0.001$. (n=6). Scale bar, 100 μm.

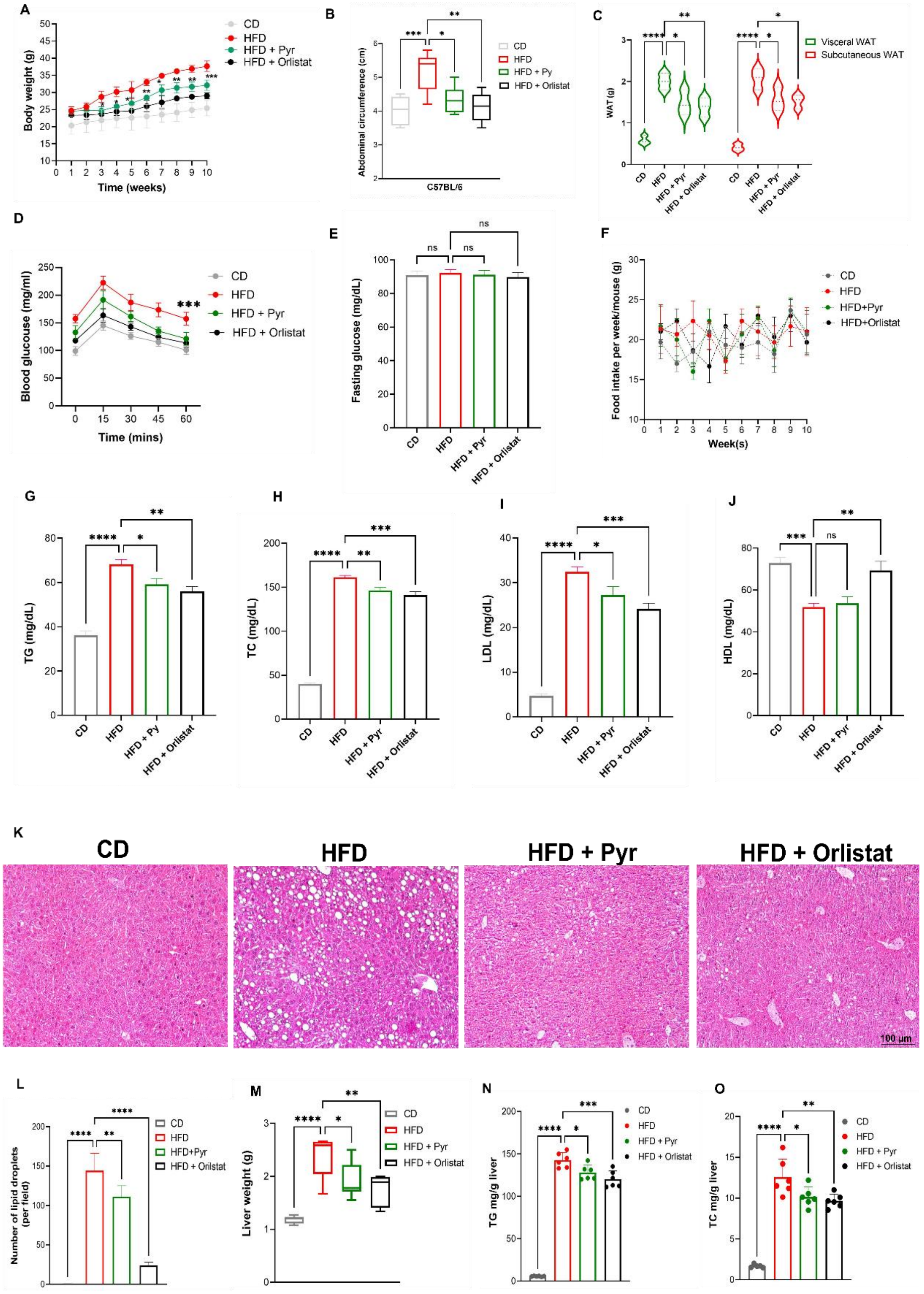


**Figure S3: Effect of pyruvate on HFD-induced obesity-induced metabolic dysregulation and hepatic steatosis in female mice.** C57BL/6 female mice were fed a high-fat diet (HFD) and chow diet (CD) for 10 weeks ± intervention with 1% pyruvate in drinking water. The experimental groups are mice fed on chow diet (CD), High-fat diet (HFD), HFD receiving pyruvate (HFD + Pyr), and HFD receiving orlistat (HFD + Orlistat).

**A.** Weight gain in female mice.

**B.** Abdominal circumference at the end of the experimental period.

**C.** Comparison between adipose tissue (vWAT and sWAT) weight.

**D.** Blood glucose level measured by OGTT after overnight fasting in the indicated experimental groups.

**E.** Fasting blood glucose level comparing different experimental groups.

**F.** Average food consumption per week.

**G-J**. Serum lipid parameters of indicated experimental group G. Triglycerides (TGs) level H. Total Cholesterol (TC) I. Low-density lipoprotein cholesterol (LDL) J. High-density lipoprotein cholesterol (HDL).

**K-L.** K. Hematoxylin and eosin-stained liver sections representing mice from each experimental group. L. Quantification of the lipid droplets (steatosis) for each group.

**M.** Weight of liver tissue.

**N.** Hepatic triglyceride level

**O.** Hepatic total cholesterol level

Data are mean ± SEM; * $p < 0.05$, ** $p < 0.01$, *** $p < 0.001$. (n=6). Scale bar, 100 µm.

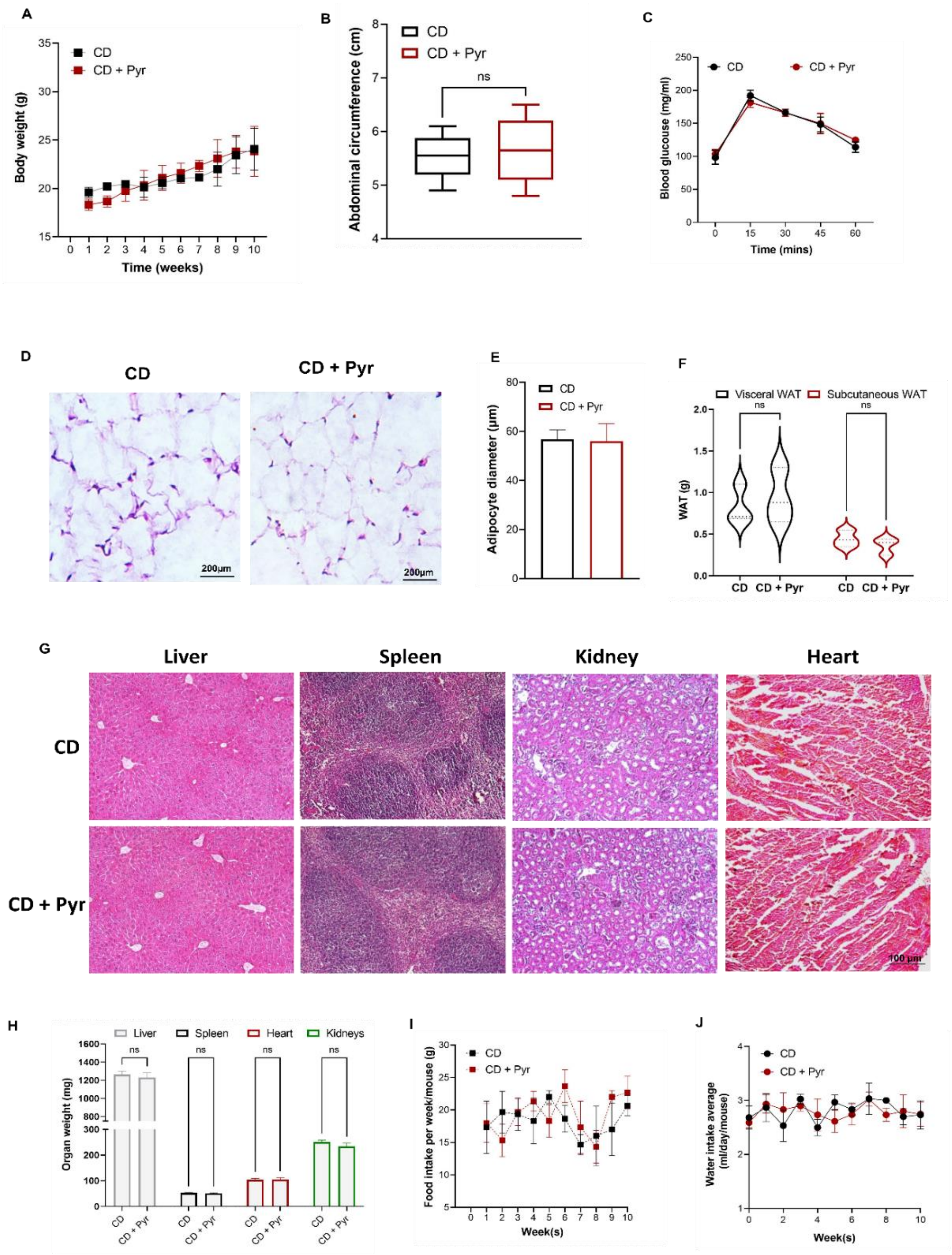


**Figure S4: Assessment of the morphologic effects of systemic pyruvate administration on CD-fed mouse tissues** **A.** Body weight gain of chow diet-fed mice over 10 weeks with (CD + Pyr) or without (CD) pyruvate intervention

**B.** Abdominal circumference at the end of the experimental period.

**C.** Blood glucose level measured by OGTT after overnight fasting in the indicated experimental groups.

**D-E.** D. Representative microphotographs of hematoxylin-eosin stained vWAT sections. E. Histological quantification for adipocyte diameter, using Image J software.

**F.** Comparison between adipose tissue (vWAT and sWAT) weight.

**G.** Representative microphotographs of hematoxylin and eosin stained indicated tissues (liver, spleen, kidney, and heart) harvested from mice following a chow diet feeding for 10 weeks with or without pyruvate supplementation in drinking water for morphometric analysis.

**H.** Organ weights

**I.** Average food consumption per week.

Data are mean ± SEM; * $p < 0.05$, ** $p < 0.01$, *** $p < 0.001$. (n=6). Scale bar, 100 μm.

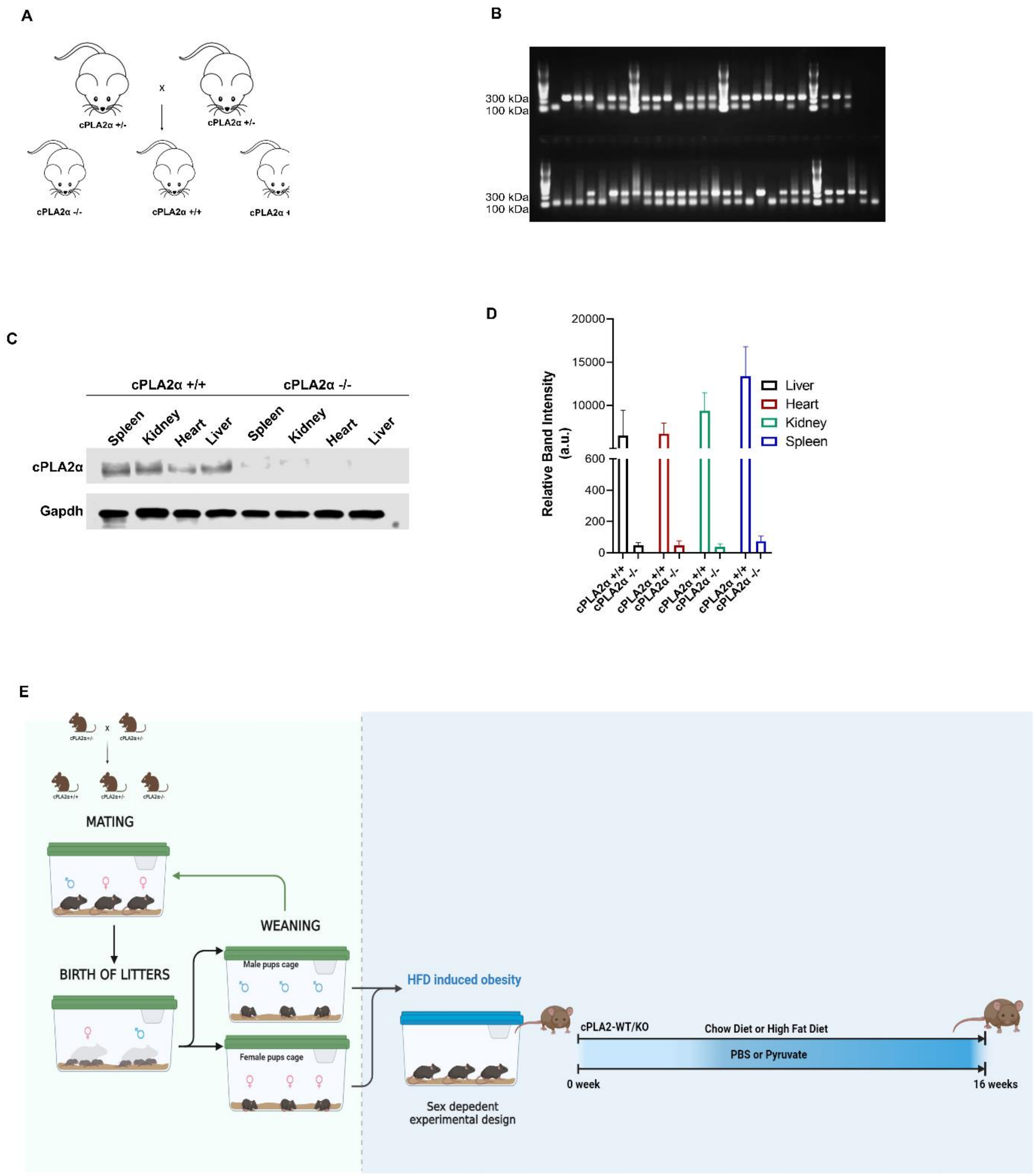


**Figure S5: Generation of global cPLA2 knockout mice**

**A.** Illustration for the breeding strategy to obtain homozygous WT (cPLA2 +/+) and homozygous KO (cPLA2 -/-) by mating cPLA2 heterozygous (cPLA2 +/-) mice.

**B**. Genotyping confirming the 100 KDa bands for the homozygous WT band, 300 kDa band for the homozygous KO band, and both bands for a heterozygous strain.

**C.** Western blot using lysates from different organs of cPLA2 WT and cPLA2 KO mice to ensure global ablation of the cPLA2 gene.

**D.** Quantification of the band intensities of panel C.

**E**. Schematic representation of the inducing HFD obesity in cPLA2 KO age-matched male mice and outlining treatment and study period.

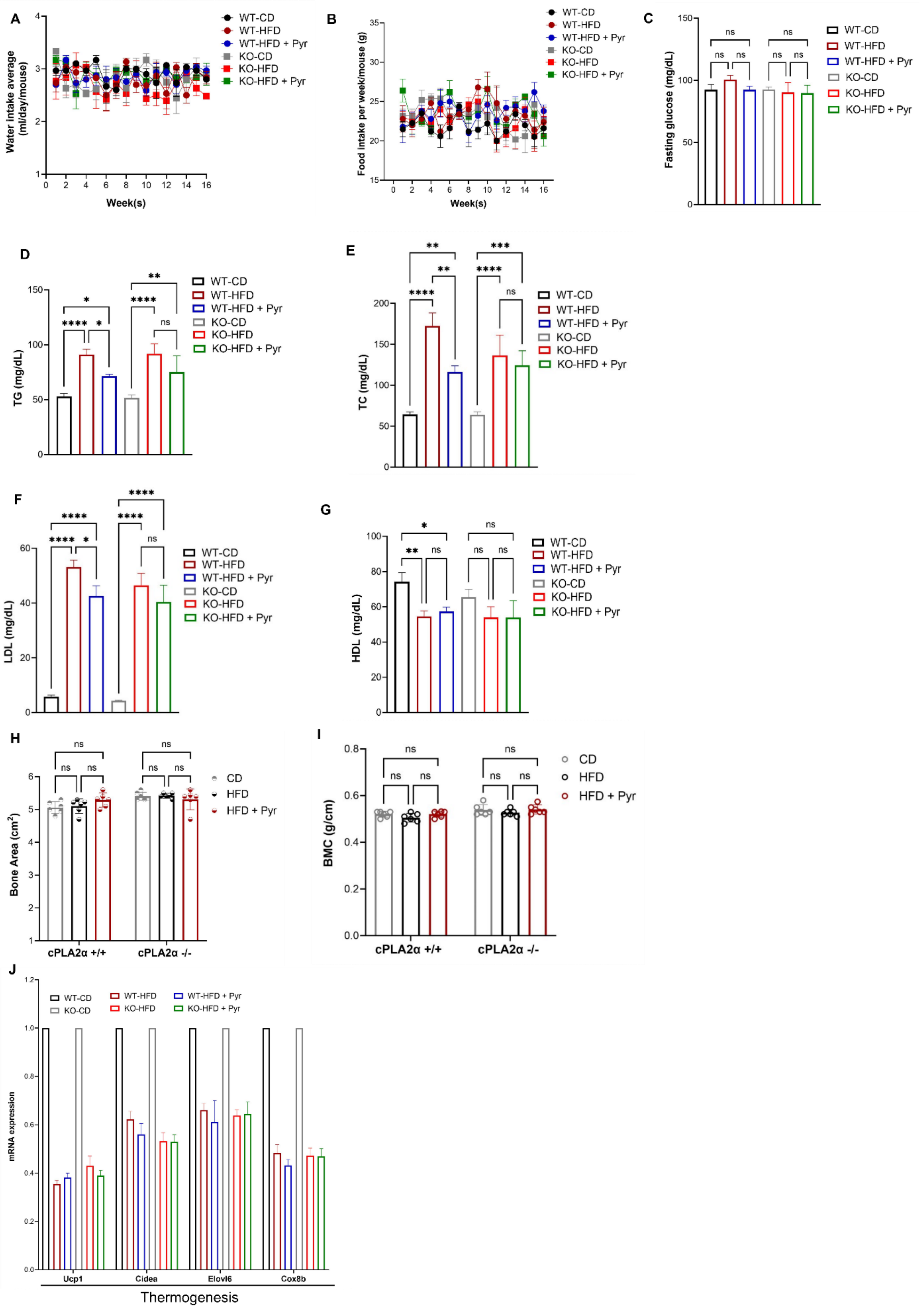


**Figure S6: cPLA2 deletion incapacitated pyruvate's ability to rescue metabolic dysfunction.**

**cPLA2 WT and cPLA2 KO mice were fed a CD or HFD for 16 weeks with or without pyruvate intervention in drinking water for the assessment of the following:**

**A.** Water consumption during the experimental period with or without intervention with 1% pyruvate in drinking water indicated a lack of aversion to pyruvate-containing water.

**B.** Average food consumption per week.

**C.** Fasting blood glucose level comparing different experimental groups.

**D-G.** Serum lipid parameters of the indicated experimental groups. D. Triglycerides level E. Total Cholesterol (TC) F. Low-density lipoprotein cholesterol (LDL) G. High-density lipoprotein cholesterol (HDL).

**H-I.** Bone indices from DXA analysis. H. Bone area and I. Bone mineral content (BMC).

**J.** Quantitative RT-PCR of the mRNA levels of markers for thermogenesis in vWAT from the indicated groups. The expression of WT and KO CD-fed mice, normalized against *Gapdh*, is regarded as 1.

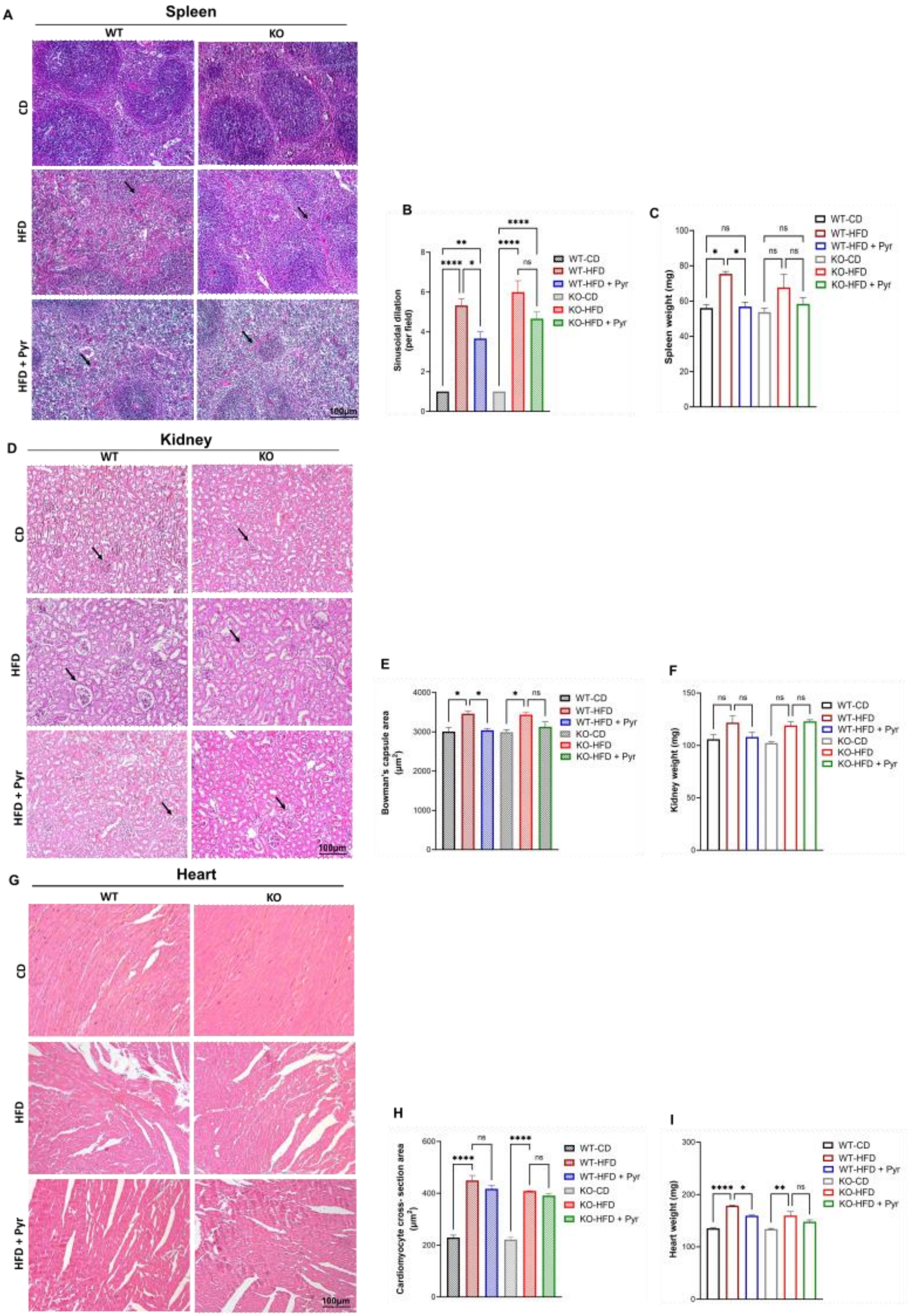
A
Spleen
WT
KO
CD
HFD
HFD + Pyr
100µm
B
Sinusoidal dilation (per field)
WT-CD
WT-HFD
WT-HFD + Pyr
KO-CD
KO-HFD
KO-HFD + Pyr
C
Spleen weight (mg)
D
Kidney
E
Bowman's capsule area (µm²)
F
Kidney weight (mg)
G
Heart
H
Cardiomyocyte cross- section area (µm²)
I
Heart weight (mg)

**Figure S7: cPLA2 deletion prevents pyruvate-mediated inhibition of ectopic fat deposition in vital organs.**

Representative microphotographs of hematoxylin and eosin stained indicated tissues harvested from cPLA2 WT and KO mice on a CD or HFD feeding for 16 weeks with or without pyruvate supplementation in drinking water.

**A-C**. A. Spleen sections with black arrows indicating the white and red pulp ratio. B. Quantification of sinusoidal dilation in each group and C. Spleen weight

**D-F**. D. Kidney sections with black arrows indicating the bowman's capsule space. E. Quantification of bowman's area in kidney sections from different experimental groups. F. Kidney weight

**G-I.** Heart sections with black arrows indicating cardiomyocytes. H. Quantification of cardiomyocyte area and I. Weight of heart tissue.

| **Primer symbol** | **Forward (5′ - 3′)** | **Reverse (5′ - 3′)** |
|---|---|---|
| IL-1β | TGGGCCTCAAAGGAAAGAAT | CAGGCTTGTGCTCTGCTTGT |
| IL-6 | CCTCTGGTCTTCTGGAGTACC | ACTCCTTCTGTGACTCCAGC |
| TNF-α | CCAGACCCTCACACTCAGATC | CACTTGGTGGTTTGCTACGAC |
| PPARγ | TGCCAGTTTCGATCCGTAGA | AGGAGCTGTCATTAGGGACATC |
| C/EBPα | CAAGAACAGCAACGAGTACCG- | TCACGCCTTTCATAACACATTCC |
| Fabp4 | AAGGTGAAGAGCATCATAACCCT | TCACGCCTTTCATAACACATTCC |
| Srebp-1c | GGAGCCATGGATTGCACATT | CTGAGTGTTTCCTGGAAGG |
| Scd1 | TTCCCTCCTGCAAGCTCTAC | CAGAGCGCTGGTCATGTAGT |
| Pnpla2 | GAGCCCCGGGGTGGAACAAGAT | AAAAGGTGGTGGGCAGGAGTAAGG |
| Elovl6 | CCAGAGCTGGCAGGTTTTACTA | CGGAGTCGCTACGTGTTCTCTA |
| UCP-1 | GTGAACCCGACAACTTCCGAA | TGCCAGGCAAGCTGAAACTC |
| Cidea | TGCTCTTCTGTATCGCCCAGT | GCCGTGTTAAGGAATCTGCTG |
| Cox8b | TGTGGGGATCTCAGCCATAGT | AGTGGGCTAAGACCCATCCTG |
| Arg1 | AGACCACAGTCTGGCAGTTG | CCACCCAAATGACACATAGG |
| Cd163 | GGGTCATTCAGAGGCACACTG | GCTGGCTGTCCTGTCAAGGCT |
| Gapdh | TGCACCACCAACTGCTTAGC | GGCATGGACTGTGGTCATGAG |
| β-actin | CCAGTTGGTAACAATGCCATGT | GGCTGTATTCCCCTCCATCG |

**Table S1:** List of mouse-specific primer sequences used for real-time PCR
